# Sequence-Specific Reduction of Interlimb Accuracy Asymmetry Reveals Preserved Motor Learning Dynamics in Chronic Stroke: Insights from Lesion-Aware fMRI

**DOI:** 10.64898/2026.08.19.745363

**Authors:** Kirstin-Friederike Heise, Patricia Finetto, Patrick A. McConnell, Christian Finetto, Femke Kiekens, Sarah E. Humphries, Seth T. Stalcup, Viswanathan Ramakrishnan

## Abstract

**Background:** People with chronic stroke retain the capacity to learn new motor skills, yet how preserved motor learning is expressed during practice remains incompletely understood. Sequence learning provides a useful model for examining these within-session learning dynamics and their neural basis after stroke.

**Objective:** To characterize a temporally resolved behavioral phenotype of motor sequence learning in chronic stroke and establish its neural context using task-based functional MRI (fMRI).

**Methods:** Twenty-four individuals with chronic stroke and 14 neurologically healthy controls performed a bimanual force-tracking sequence-learning task during functional MRI. Performance convergence was defined as the sequence-specific reduction in the accuracy difference between the paretic and less-affected hands across practice. Neural activity was evaluated using whole-brain, region-of-interest, and functional-connectivity analyses following preprocessing tailored to structurally heterogeneous stroke lesions.

**Results:** Stroke participants demonstrated significant performance convergence despite persistent motor impairment, indicating preserved expression of sequence learning during practice that was not detected by conventional behavioral measures. Lesion-aware fMRI identified robust task-related activation and preserved stage-dependent modulation within cerebellar, premotor, and striatal learning networks, together with reduced bilateral putaminal activity after stroke. However, preregistered analyses found no reproducible associations between individual differences in performance convergence and learning-related activation or functional connectivity.

**Conclusions:** Performance convergence provides a sensitive, temporally resolved behavioral phenotype of preserved motor sequence learning in chronic stroke that complements conventional endpoint measures. Together, performance convergence and task-based functional MRI provide a framework for investigating individual differences in motor learning capacity and their implications for rehabilitation responsiveness.

**ClinicalTrials.gov:** NCT05511467.

## Introduction

Experience-dependent plasticity enables the nervous system to modify behavior in response to changing environmental demands and is a principal biological substrate supporting recovery after stroke^1–3^. Although considerable evidence demonstrates that people with chronic stroke retain the capacity to acquire new motor skills^4–6^, motor learning after stroke is commonly summarized using aggregate outcomes such as overall accuracy, total improvement, or an estimated learning rate^7^. Such summaries do not resolve the trajectory through which learning unfolds and may overlook transient changes or differences between limbs that emerge during practice^8,9^. Better characterization of within-session behavioral dynamics is needed to determine whether preserved components of motor learning remain hidden when learning is summarized by endpoint performance.

Motor sequence learning provides a useful model because learning unfolds through temporally structured stages that can be resolved within a single practice session^8,10,11^. Early learning recruits distributed sensorimotor, associative, and executive systems^9,10,12,13^, whereas continued practice is accompanied by reorganization of corticostriatal, cerebellar, premotor, and parietal networks as sequence knowledge improves^10,11,14^, enabling increasingly predictive movement planning and more efficient motor control^15,16^. Characterizing these dynamics after stroke therefore requires examining how performance changes throughout practice rather than simply whether learning occurs.

Stroke introduces additional complexity because motor learning occurs in the context of heterogeneous structural injury and altered large-scale network organization^17–19^. Rather than simply reducing motor capacity, focal lesions alter activity and connectivity within distributed motor-learning networks that support the acquisition and refinement of skilled behavior^5,20,21^. Aggregate performance measures may therefore fail to capture preserved learning-related processes that become apparent only when behavioral trajectories are examined over time. We reasoned that, as sequence knowledge increasingly supports advance planning during practice, the paretic hand should progressively benefit from this predictive information, producing a sequence-specific reduction in interlimb accuracy asymmetry relative to random practice. We define this behavioral trajectory as *performance convergence*, reflecting increasing expression of predictive sequence knowledge by the paretic hand, and hypothesized that it provides a sensitive marker of preserved motor learning dynamics after stroke.

Establishing the neural context of such behavioral dynamics presents a methodological challenge. Task-based functional MRI provides a noninvasive means of characterizing learning-related neural activity across distributed motor networks^5,20^. However, focal lesions bias spatial normalization and produce heterogeneous functional coverage, complicating both individual- and group-level inference. We therefore combined explicit lesion masking, voxelwise coverage assessment, and lesion-load analysis to support robust whole-brain and regionally constrained analyses. Together, these behavioral and methodological approaches provide a framework for investigating the neural systems associated with individualized motor learning trajectories after stroke.

In the present study, we combined a bimanual force-tracking sequence-learning paradigm with lesion-aware task fMRI to investigate within-session motor learning in chronic stroke. First, we tested whether chronic stroke preserves performance convergence, operationalized as a sequence-specific reduction in interlimb accuracy asymmetry. Second, we characterized learning-related brain activity using a lesion-aware task-fMRI framework designed to accommodate substantial lesion heterogeneity. Third, we tested whether individual differences in performance convergence were associated with corresponding changes in task-related brain activity and functional connectivity.

By integrating temporally resolved behavioral analysis with lesion-aware neuroimaging, this study evaluates performance convergence as a candidate behavioral phenotype of preserved motor learning after stroke.

## Methods

Comprehensive methods, statistical models, quality-control procedures, and additional analyses are provided in the Supplementary Material (**Supplementary Methods, Tables S1–S2; Fig. S1**).

### Participants and Baseline Characterization

Individuals with chronic stroke (≥6 months) and neurologically healthy volunteers were prospectively recruited for this preregistered observational study (ClinicalTrials.gov: NCT05511467; participant flow, **Fig. S1**). Following exclusions, the final sample comprised 24 stroke and 14 control participants. The study was approved by the Institutional Review Board of the Medical University of South Carolina (Pro00116626) and conducted in accordance with the Declaration of Helsinki. All participants provided written informed consent before participation.

All participants underwent standardized neurological, sensorimotor, neurocognitive, and behavioral characterization before the experimental session. Stroke participants additionally completed standardized measures of impairment and upper-extremity function (NIHSS^22,23^, UE-FMA^24^, ARAT^25,26^, 9HPT^27,28^, SIS-3.0^29,30^), while all participants completed assessments of handedness^31^, cognition^32^, mood^33^, sleep^34–36^, medication use, and potential state-dependent confounds.

Baseline demographic, clinical, and behavioral characteristics were summarized separately for the stroke and control groups (**Table 1**). Continuous variables were compared using Welch’s t-tests or Mann–Whitney U tests, as appropriate, and categorical variables using Pearson’s χ² or Fisher’s exact tests.

**Table 1.**
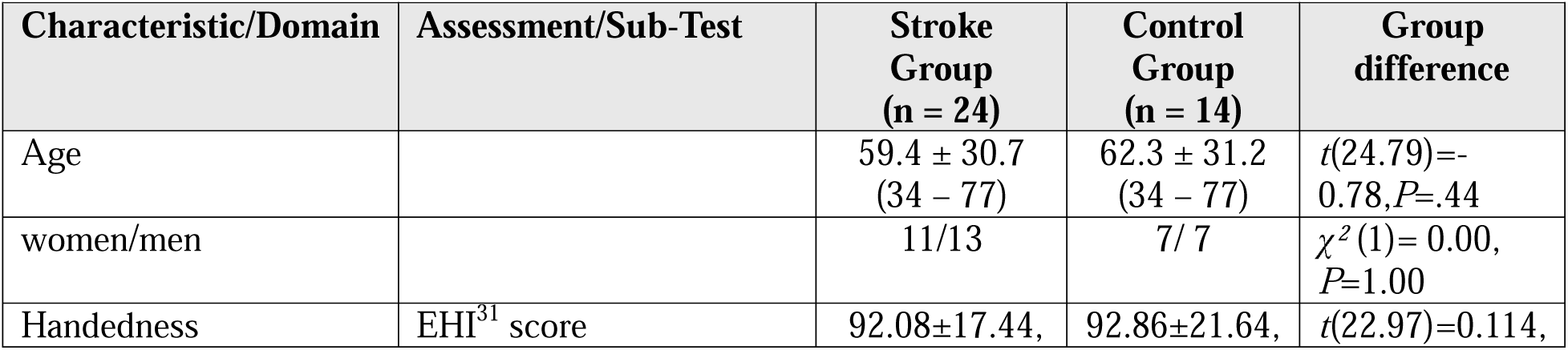
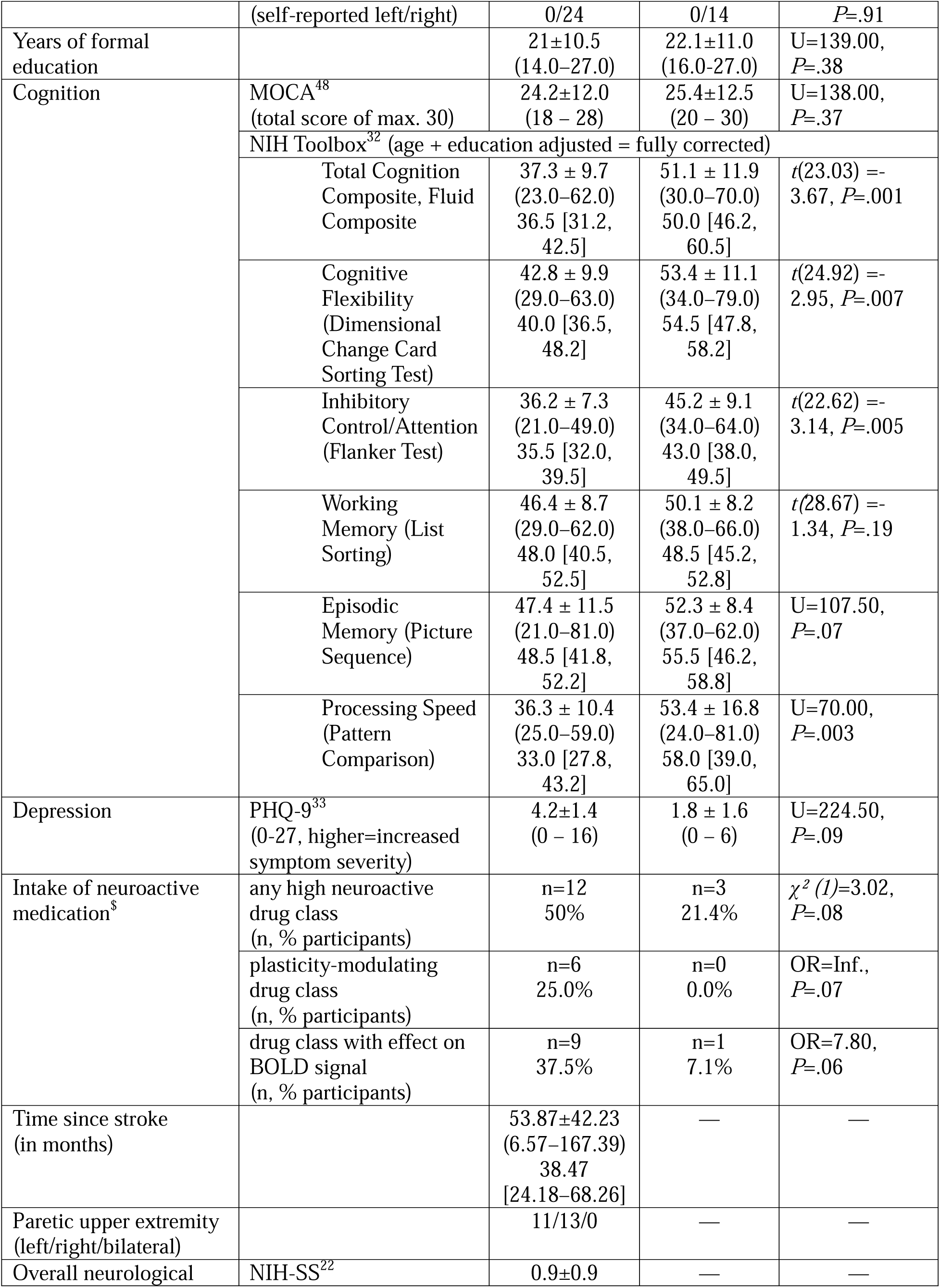
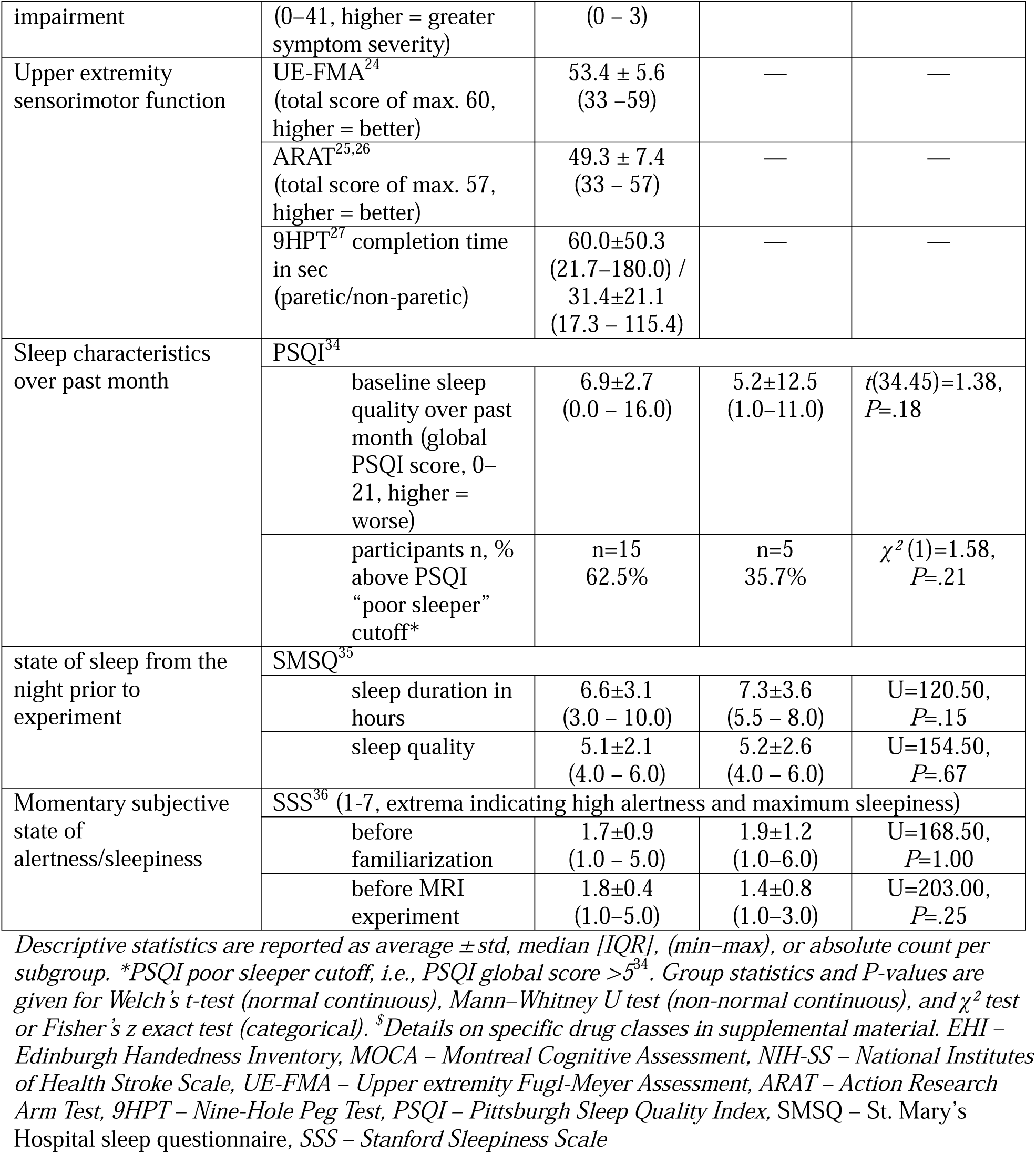
Sample Characteristics.

### Study Design and Experimental Procedures

An overview of the experimental design and bimanual force-tracking paradigm is shown in **Fig. 1**. Participants completed baseline characterization, a practice session before MRI, and a task-fMRI experiment comprising sequence (SEQ) and random (RND) force-tracking conditions across 3–4 sessions spread over 1–2 weeks.

**Fig. 1.**
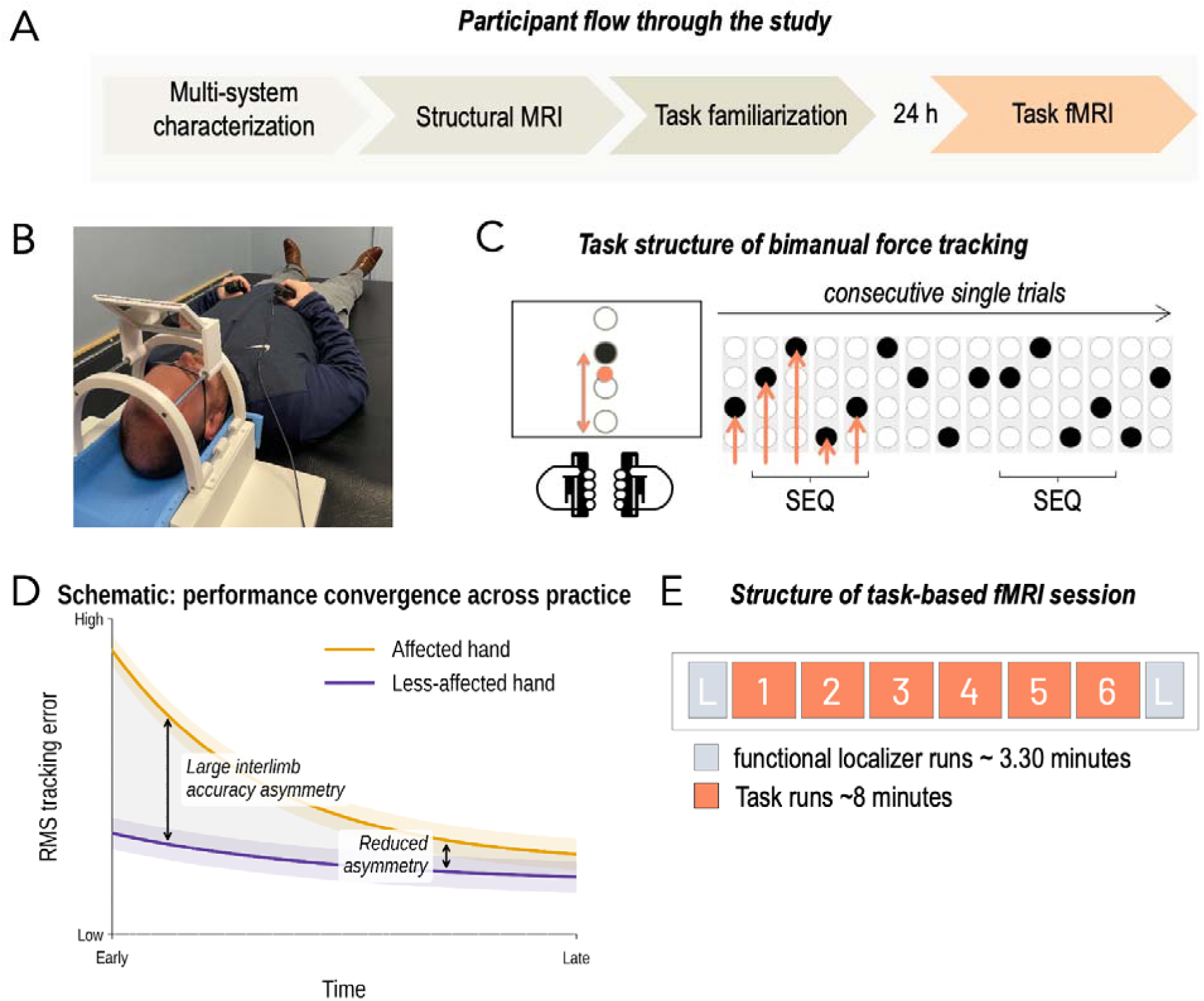
Study design and conceptual framework for performance convergence. **A)** Participants completed standardized neurocognitive and sensorimotor assessments followed by structural and functional MRI. **B)** Approximately 24 hours before task-based fMRI, participants completed a practice session in a setting that reproduced the in-scanner task presentation and response setup; one of the authors shown. **C)** During the bimanual force-tracking task, one of four target force levels, scaled to 4%–7% of each hand’s maximum voluntary contraction, was indicated by a visual target (black circle). Participants synchronously modulated grip force with both hands to move a single response cursor (orange), representing the combined bilateral force output, to the target, maintained the required force until target offset, and then released both transducers. Although visual feedback was based on the combined bilateral force signal, grip force from each hand was recorded separately to derive hand-specific tracking accuracy and interlimb accuracy asymmetry. Target-presentation lasted 2s and intertrial intervals were jittered between 1.5–2.5 s. **D)** Conceptual illustration of **performance convergence**, defined as the progressive reduction in interlimb accuracy asymmetry during sequence learning as tracking accuracy of the paretic hand progressively approached that of the less-affected hand. **E)** Task-based fMRI comprised six experimental runs (∼8 min each) including counterbalanced blocks of sequence (SEQ) and random (RND) force-tracking. RND-only functional localizer runs (L) were acquired immediately before and after the experimental runs. These runs were used to quantify condition-general change across the scanning session and characterize force-level-related activation.

### Sequence learning implemented with the bimanual force-tracking (BFT)

During the BFT task, participants continuously modulated bilateral grip force to match visually presented target forces while grip force was recorded at 1000 Hz. Target forces were scaled to 4– 7% of each participant’s maximum voluntary contraction (re-adjusted before each run). Six experimental runs alternated blocks with exclusively sequential (SEQ) or random (RND) trials, and RND-only localizer runs acquired immediately before and after the experiment. Primary learning analyses used the experimental runs; localizers quantified condition-general change.

#### Primary behavioral outcome: interlimb accuracy asymmetry and performance convergence

The primary behavioral analysis tested whether chronic stroke alters the within-session trajectory of sequence-specific change in the relative accuracy of the two hands. For each trial, *interlimb accuracy asymmetry* was defined as the difference in root-mean-square (RMS) force-tracking error between the paretic and less-affected hands (controls: non-dominant minus dominant hand). Performance convergence was operationalized as the sequence-specific reduction in interlimb accuracy asymmetry during practice. Trial-level asymmetry was analyzed using a Group × Condition × Run linear mixed-effects model in which Run was modeled as a continuous, mean-centered predictor, with a by-subject random intercept and a random-effects structure selected as detailed in the **Supplementary Methods**; subject-specific performance-convergence slopes were estimated from the SEQ runs for subsequent brain–behavior analyses. Subject-specific performance-convergence slopes derived from SEQ runs served as brain–behavior covariates after winsorization (median ±3 MAD). RND-only localizer analyses quantified condition-general change. Significant interactions were further decomposed using estimated marginal means and slope contrasts (R package emmeans), Holm-corrected within each family of comparisons. Complete model statistics and follow-up contrasts are reported in **Tables S3–S5**.

#### Secondary behavioral analyses

Secondary analyses modeled trial-level accuracy and reaction time (RT) using linear mixed-effects models and compared pre-versus post-task localizer performance to quantify condition-general change. Model outputs are summarized in **Tables S4–S5**.

### MRI acquisition and neuroimaging analyses

#### MRI data acquisition and lesion-aware preprocessing

MRI data were acquired on a 3T Siemens Prisma fit system (Siemens Healthineers, Erlangen, Germany) using a 32-channel head coil. High-resolution structural images were obtained using a T1-weighted 3D MPRAGE sequence (TR = 2.3 s, TE = 2.26 ms, TI = 900 ms, flip angle = 8°, 1- mm isotropic resolution, GRAPPA factor = 2). Whole-brain BOLD images were acquired using a T2*-weighted gradient-echo echo-planar imaging sequence (TR = 1.5 s, TE = 30 ms, flip angle = 65°, multiband factor = 3, in-plane acceleration = 2, 2-mm isotropic resolution, phase-encoding direction j−). Dual-echo GRE field maps were acquired for distortion correction and T2-weighted FLAIR images for lesion characterization. Imaging data were converted to BIDS^37^ format using dcm2niix^38^ and CuBIDS^39^.

Functional and anatomical data were initially preprocessed using fMRIPrep^40^ with its standard anatomical and functional workflow. Because focal lesions degrade automated cortical surface reconstruction, FreeSurfer surfaces were generated externally (FreeSurfer 7.4.1) and supplied to fMRIPrep as precomputed input; all stroke reconstructions were visually inspected and manually edited to correct reconstruction failures in and around the lesion before use. Preprocessed functional images were resampled to MNI152NLin2009cAsym space at 2-mm isotropic resolution using fMRIPrep’s anatomical-to-template transform, which was retained for all participants; the manually segmented lesions were not used to modify spatial normalization.

To account for heterogeneous structural lesions, stroke lesions were manually segmented on each participant’s structural images using ITK-SNAP^41^, visually reviewed and approved by a board-certified neuroradiologist. The challenge of spatially normalizing lesioned brains is well recognized^42–44^; rather than modifying the registration to account for the lesion, we reused fMRIPrep’s anatomical-to-template transform to resample the functional images, brain masks, and lesion masks into MNI space, and incorporated lesion information at the masking, coverage, and analysis stages — the sense in which the pipeline is lesion-aware. This produced spatially congruent data across participants. Additional preprocessing details are provided in the **Supplementary Methods**. For each stroke participant, an explicit first-level analysis mask was generated by excluding the normalized lesion mask from the corresponding brain mask. Normalized lesion masks were additionally combined to characterize lesion distribution and group-level lesion frequency.

#### First-level activation analyses

Task-fMRI analyses were conducted in SPM25^45,46^ using the six experimental BFT runs; the pre- and post-task localizer runs were excluded from these analyses. Preprocessed functional images were smoothed using a 6-mm full-width-at-half-maximum Gaussian kernel. Subject-specific explicit masks combined anatomically valid non-lesioned tissue with functional signal-coverage criteria to restrict model estimation to reliably sampled voxels.

A separate first-level general linear model was estimated for each participant, with one session per run. Sequential (SEQ) and random (RND) task conditions were modeled using the canonical hemodynamic response function and temporal derivative. Run-specific nuisance regressors included motion parameters, their temporal derivatives, and outlier regressors. Low-frequency signal drift was removed using a 128-s high-pass filter, and temporal autocorrelation was modeled using SPM’s first-order autoregressive model.

First-level contrasts quantified SEQ and RND task activity, sequence-specific activity (SEQ−RND), early-to-late change, and early-to-late change in the sequence-specific effect. These participant-level contrast images were entered into group-level random-effects analyses.

#### Group-level activation analyses

Group-level analyses were conducted separately for the stroke and control cohorts. Because lesion location and functional coverage varied across stroke participants, contrast-specific explicit group masks were generated from the proportion of participants with valid contrast estimates at each voxel. Coverage thresholds were set at ≥50% of participants in the stroke group and ≥80% in the control group and were intersected with a canonical brain-tissue mask.

Whole-brain analyses characterized task-related and learning-stage activity and tested associations between performance-convergence slopes and early-to-late sequence-specific activation. Preregistered analyses used the winsorized behavioral covariate in stroke participants; raw covariates and control analyses served as sensitivity and specificity analyses.

#### Lesion-load analyses

To evaluate whether structural lesion burden modified the association between performance convergence and learning-related neural activity, whole-brain and bilateral primary motor cortex lesion-load indices were derived from the subject-specific analysis masks. These indices quantified the proportion of expected tissue coverage lost within each anatomical domain and were corrected for generic coverage and registration variability using the corresponding median values in control participants.

Stroke-only second-level regression models tested whether lesion load moderated the association between the performance-convergence slope and the sequence-specific early-to-late activation contrast. Centered performance-convergence and lesion-load terms and their interaction were entered simultaneously.

#### Region-of-interest (ROI) analyses

A priori region-of-interest analyses targeted cortical, subcortical, and cerebellar nodes implicated in visuomotor sequence learning and its stage-dependent reorganization. Anatomical ROIs included bilateral primary motor cortex, superior parietal lobule, caudate, putamen, and lateral occipital cortex. Functional ROIs targeted pre-supplementary motor area, dorsal and ventral premotor cortex, and cerebellar lobule VI.

Mean contrast estimates were extracted from each ROI after accounting for participant-specific lesion and signal coverage. Estimates were retained when at least 25% of the ROI contained valid data; analyses using a stricter 50% coverage threshold were conducted as sensitivity analyses. ROI analyses examined sequence-specific and learning-stage-dependent effects. Between-group effects were assessed using Welch’s t tests, with nonparametric tests used as sensitivity analyses, and within-group stage effects were assessed using paired comparisons.

False-discovery-rate correction was applied within each prespecified family of ROI tests. ROI definitions are listed in **Table S2**.

#### Functional-connectivity analyses

Generalized psychophysiological interaction (gPPI)^47^ analyses were used to assess task-modulated functional connectivity. The primary connectivity analysis focused on interhemispheric coupling between ipsilesional and contralesional primary motor cortex hand-knob regions, defined relative to the affected hand in stroke participants and the assigned non-dominant hand in controls. Additional seeds in pre-supplementary motor, premotor, superior-parietal, putaminal, and dorsal anterior cingulate regions were examined as secondary or exploratory analyses. Seed-specific gPPI models quantified condition-specific connectivity, sequence-specific effects (SEQ−RND), and early-to-late changes in coupling.

Second-level connectivity analyses tested task-modulated connectivity, learning-related change, associations with the performance-convergence slope, and moderation by lesion load. The pre-specified primary connectivity tests focused on the bilateral M1 seed pair in the stroke cohort. Raw-covariate and control analyses served as sensitivity and specificity analyses.

#### Statistical inference and multiple-comparison control

Confirmatory and exploratory analyses were prespecified. Five PRIMARY brain–behavior analyses comprised whole-brain activation, lesion-load moderation, and three bilateral M1 gPPI models; all remaining analyses were secondary or exploratory. Whole-brain inference used voxel-wise FWE correction (p<.05); uncorrected results (p<.001, k≥10) are reported descriptively.

## Results

Complete behavioral data were available for 24 stroke participants and 14 control participants. The groups were comparable in demographic characteristics and global cognition, although stroke participants showed lower performance on selected NIH Toolbox measures, particularly executive function and processing speed (**Table 1**). Lesion location was heterogeneous across the stroke cohort (**Fig. 2A**; **Table 2**). Imaging sample sizes varied after quality control and are reported for each analysis below.

**Fig. 2.**
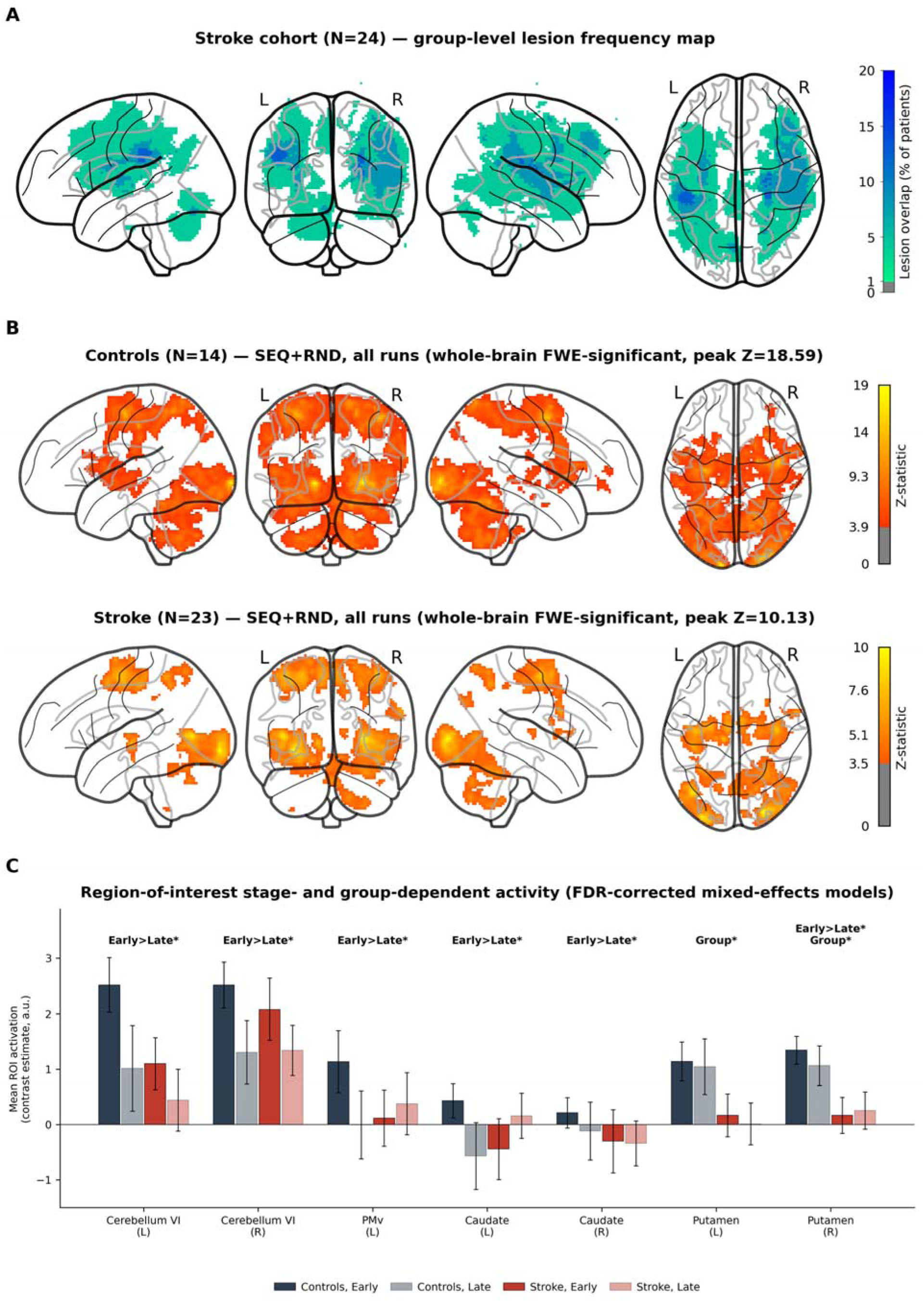
Lesion-aware task-fMRI identifies preserved learning-stage modulation and reduced putaminal activity after stroke. **A)** Group-level lesion frequency map for the stroke cohort (N = 24), displayed on the MNI152 nonlinear template using a glass-brain maximum-intensity projection derived from individual binary lesion masks. Colors indicate the percentage of participants with lesion involvement at each voxel. This panel includes all participants with successfully normalized lesion masks, including one participant excluded from the functional MRI analyses because reliable BOLD spatial normalization could not be achieved. **B)** Whole-brain one-sample activation maps for the combined task-positive contrast (SEQ + RND) in stroke and control participants, thresholded at voxel-wise family-wise-error (FWE) corrected p < .05, demonstrating robust activation within the expected visuomotor network. **C)** Region-of-interest analyses of the same visuomotor network showed early-to-late decreases in bilateral cerebellar lobule VI, left ventral premotor cortex, bilateral caudate, and right putamen (all FDR q<.05), together with lower bilateral putamen activity in stroke independent of learning stage. Bars show group means ± SEM of per-subject regional contrast estimates, averaged across task conditions. Labels above each region indicate which FDR-corrected (q<.05) mixed-effects-model effects reached significance (Results, “Region-of-interest findings”), not the raw group means plotted: “Early>Late” denotes a significant pooled early-to-late decrease across groups and task conditions, and “Group” denotes significant pooled stroke<control difference across stage and condition. Complete ROI statistics are reported in **Table S7**.

**Table 2.**
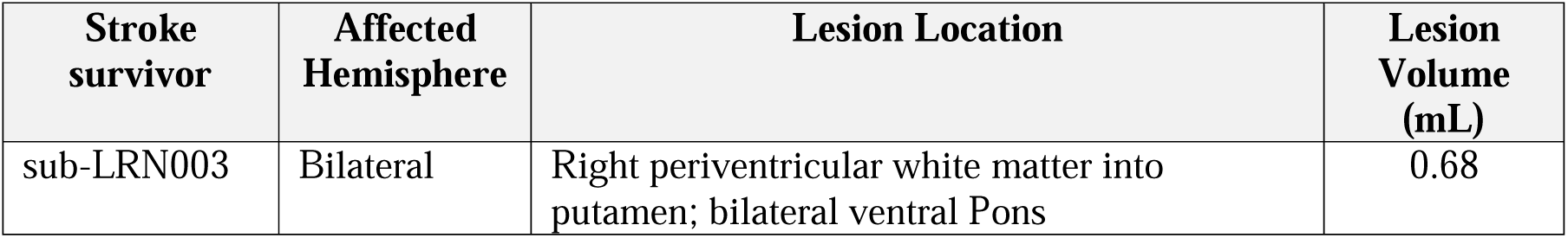
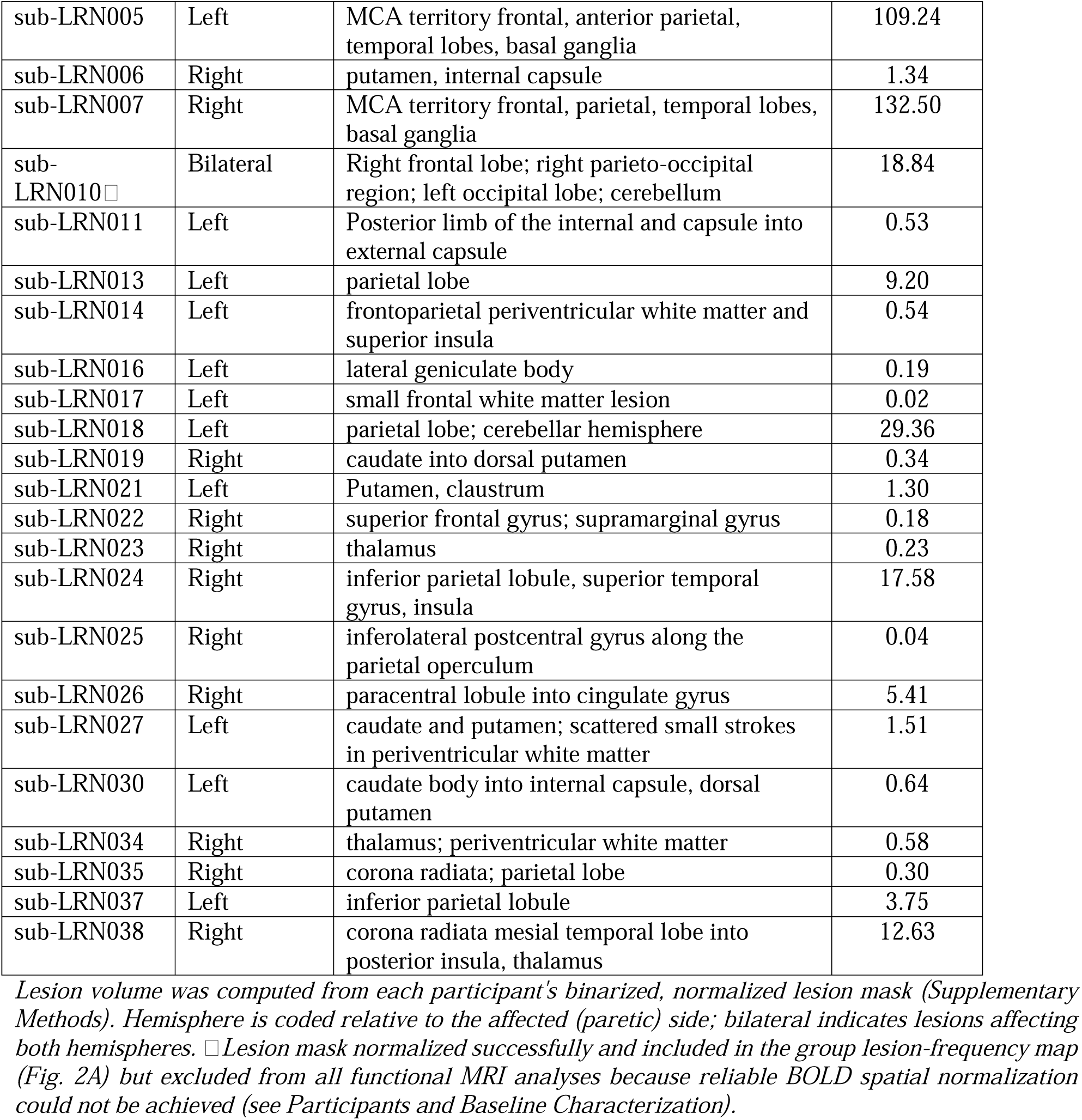
Lesion Characteristics.

| Stroke survivor | Affected Hemisphere | Lesion Location | Lesion Volume (mL) |
| --- | --- | --- | --- |
| sub-LRN003 | Bilateral | Right periventricular white matter into putamen; bilateral ventral Pons | 0.68 |
| Running title: Performance Convergence After Stroke |  |  |  |
| sub-LRN005 | Left | MCA territory frontal, anterior parietal, temporal lobes, basal ganglia | 109.24 |
| sub-LRN006 | Right | putamen, internal capsule | 1.34 |
| sub-LRN007 | Right | MCA territory frontal, parietal, temporal lobes, basal ganglia | 132.50 |
| sub-LRN010 □ | Bilateral | Right frontal lobe; right parieto-occipital region; left occipital lobe; cerebellum | 18.84 |
| sub-LRN011 | Left | Posterior limb of the internal and capsule into external capsule | 0.53 |
| sub-LRN013 | Left | parietal lobe | 9.20 |
| sub-LRN014 | Left | frontoparietal periventricular white matter and superior insula | 0.54 |
| sub-LRN016 | Left | lateral geniculate body | 0.19 |
| sub-LRN017 | Left | small frontal white matter lesion | 0.02 |
| sub-LRN018 | Left | parietal lobe; cerebellar hemisphere | 29.36 |
| sub-LRN019 | Right | caudate into dorsal putamen | 0.34 |
| sub-LRN021 | Left | Putamen, claustrum | 1.30 |
| sub-LRN022 | Right | superior frontal gyrus; supramarginal gyrus | 0.18 |
| sub-LRN023 | Right | thalamus | 0.23 |
| sub-LRN024 | Right | inferior parietal lobule, superior temporal gyrus, insula | 17.58 |
| sub-LRN025 | Right | inferolateral postcentral gyrus along the parietal operculum | 0.04 |
| sub-LRN026 | Right | paracentral lobule into cingulate gyrus | 5.41 |
| sub-LRN027 | Left | caudate and putamen; scattered small strokes in periventricular white matter | 1.51 |
| sub-LRN030 | Left | caudate body into internal capsule, dorsal putamen | 0.64 |
| sub-LRN034 | Right | thalamus; periventricular white matter | 0.58 |
| sub-LRN035 | Right | corona radiata; parietal lobe | 0.30 |
| sub-LRN037 | Left | inferior parietal lobule | 3.75 |
| sub-LRN038 | Right | corona radiata mesial temporal lobe into posterior insula, thalamus | 12.63 |

### Sequence-specific reduction in interlimb accuracy asymmetry

The primary behavioral analysis tested whether the accuracy difference between the affected and less-affected hands changed across sequence-learning practice. At initial exposure, the size of the interlimb accuracy asymmetry depended jointly on group and condition (significant Group × Condition interaction, χ²(1) = 4.36, p = .037); follow-up simple contrasts did not reach significance within either condition alone (all p ≥ .22). Critically, this imbalance changed across runs in a group- and condition-dependent manner, as indicated by a significant Group × Condition × Run interaction (χ²(1) = 5.76, p = .016; **Fig. 3**). Follow-up contrasts localized this interaction to a significantly steeper (more negative) run-slope during SEQ than RND practice within stroke participants specifically; the direct stroke-vs-control comparison of SEQ slopes did not independently reach significance. The significant three-way interaction, rather than the within-group slope contrast alone, provides the primary behavioral evidence that within-session interlimb rebalancing across practice depended jointly on group and condition. Complete model statistics are provided in **Tables S3–S5**.

**Fig. 3.**
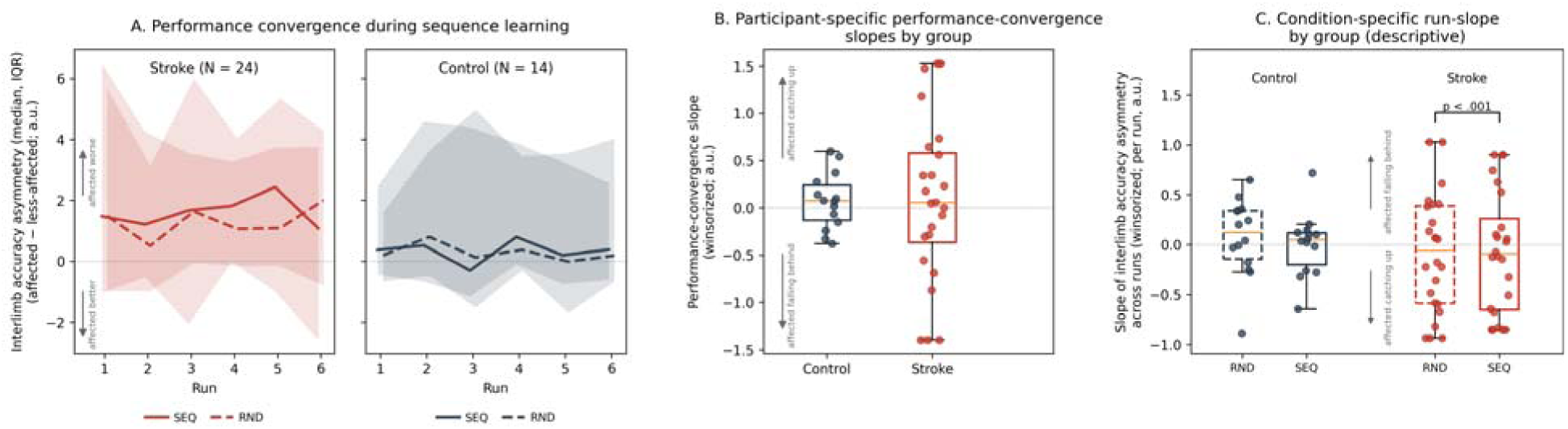
Sequence-specific performance convergence reveals preserved motor learning in chronic stroke. **A)** Interlimb accuracy asymmetry (affected minus less-affected hand) across task runs for sequential (SEQ, solid line) and random (RND, dashed line) conditions, shown separately for stroke (N = 24) and control (N = 14) participants on a shared y-axis. Lines and shaded bands show the median and interquartile range across participants at each run, reducing the influence of a small number of outlying subject-runs. Arrows and adjacent labels indicate that positive values reflect worse accuracy in the affected (or assigned non-dominant in controls) hand relative to the less-affected (or dominant) hand, and negative values reflect better accuracy. The figure provides a descriptive summary; statistical inference is based on the trial-level linear mixed-effects models reported in Supplementary Results. **B)** Participant-specific performance-convergence slopes derived from the primary mixed-effects model (winsorized; median ± 3 MAD) and used as the behavioral covariate in subsequent brain–behavior analyses, shown by group. Arrows and adjacent labels indicate that positive values reflect the affected hand catching up to the less-affected hand across runs, and negative values reflect it falling further behind. **C)** Condition-specific run-slopes of interlimb accuracy asymmetry, computed per participant via simple linear regression across runs separately for SEQ and RND trials (winsorized; median ± 3 MAD), shown by group and condition. Arrows and adjacent labels follow the same convention as panel A. Bracket indicates the only model-based follow-up contrast that remained significant after Holm correction: within stroke participants, the SEQ run-slope was significantly more negative than the RND run-slope (p < .001); no other pairwise contrast was significant. Complete mixed-effects models and follow-up contrasts are reported in **Tables S3–S5**.

To determine whether this narrowing was specific to sequence learning, an analogous analysis examined changes during random-only force tracking during pre- and post-localizer runs. The Group × Timepoint interaction was also significant for RND performance (χ²(1) = 7.47, p = .006). Follow-up contrasts showed this reduction was significant within stroke participants (pre-to-post decrease, p < .001) but not within control participants (p = .691), indicating that this condition-general component of the reduction was itself concentrated in the stroke group rather than reflecting a generic session effect shared by both groups. The stronger three-way interaction in the primary model nevertheless showed that the trajectory of interlimb rebalancing differed across task conditions and groups. Accordingly, throughout the remainder of this manuscript, the term performance convergence refers specifically to the sequence-specific reduction in interlimb accuracy asymmetry beyond the condition-general changes observed during random practice. A participant-specific performance-convergence slope derived from the SEQ model was therefore used as the behavioral covariate in subsequent brain–behavior analyses.

#### Secondary characterization of sequence learning and session-related change

Secondary analyses examined trial-level accuracy and reaction time across both hands to determine whether the primary interlimb effect was also evident in conventional performance measures. As expected, the affected hand showed poorer accuracy and slower responses than the less-affected hand (accuracy: χ²(1) = 96.56, p < .001; reaction time: χ²(1) = 45.65, p < .001).

For accuracy, a significant Group × Condition interaction (χ²(1) = 12.99, p < .001) indicated that the sequence-vs-random accuracy difference depended on group; follow-up simple group contrasts within each condition did not individually reach significance (RND: p = .195; SEQ: p = .077, Table S5). This interaction remained significant when learning-related changes across runs were included in the model (χ²(1) = 13.52, p < .001). However, the Group × Condition × Run interaction was not significant for trial-level accuracy (χ²(1) = 0.08, p = .78), indicating that the group difference in overall sequence benefit did not itself change across practice. Reaction time showed neither a significant Group × Condition interaction (χ²(1) = 2.70, p = .10) nor a Group × Condition × Run interaction (χ²(1) = 1.18, p = .28).

Condition-general changes across the scanning session were assessed using the RND-only localizer runs acquired before and after the experimental task. Accuracy declined from pre- to post-task assessment (χ²(1) = 9.84, p = .002), but this change did not differ significantly between groups (Group × Timepoint: χ²(1) = 2.47, p = .12). Reaction time also changed across the session (χ²(1) = 14.21, p < .001), with a significant Group × Timepoint interaction (χ²(1) = 7.33, p = .007); follow-up contrasts showed this reflected significant slowing from pre- to post-task assessment in control participants (p < .001), with no significant change in stroke participants (p = .746).

In summary, performance convergence emerged as a temporally resolved behavioral phenotype of motor sequence learning that was not captured by conventional endpoint measures. The RND and localizer analyses further demonstrated that this effect reflected sequence-specific learning rather than condition-general session effects.

### Neuroimaging results

Whole-brain activation analyses included 23 stroke participants and 14 control participants after quality control. One additional stroke participant was excluded because spatial normalization failed in the presence of multiple large lesions. Head-motion, normalization, and regional-coverage metrics were acceptable in the retained sample.

#### Task-related and learning-related activation

Because lesion location produced heterogeneous voxelwise coverage in the stroke cohort, all group-level analyses were conducted using cohort- and contrast-specific explicit masks. These masks retained broad frontal, parietal, visual, and sensorimotor coverage while restricting inference to voxels sampled in a sufficient proportion of participants.

Whole-brain one-sample analyses demonstrated robust task-related activation in both groups. SEQ and RND conditions, as well as their conjunction, produced anatomically expected family-wise-error-corrected responses in visual, sensorimotor, and premotor regions (**Fig. 2B**), confirming adequate sensitivity for detecting the principal task-evoked response.

In contrast, no learning-related whole-brain effects survived family-wise-error correction for the early-to-late, early SEQ−RND, or late SEQ−RND contrasts in either group. A small cluster observed in controls under an alternative smoothing parameter did not replicate across analysis specifications and was not interpreted further. Thus, although the pipeline detected robust task-related activity, the present sample did not provide stable whole-brain evidence for more subtle learning-stage effects.

#### Region-of-interest findings

A priori ROI analyses examined stage- and group-dependent activity within the visuomotor sequence-learning network. Of 33 tests across 11 ROIs and three effect families, eight survived false-discovery-rate correction.

Activity decreased from early to late practice in bilateral cerebellar lobule VI, left ventral premotor cortex, bilateral caudate, and right putamen (all FDR-corrected q < .05, **Fig. 2C**). Independent of learning stage, bilateral putamen activity was lower in stroke participants than in controls (left: F(1,37) = 9.12, p = .005, q = .031; right: F(1,37) = 8.62, p = .006, q = .031). No Group × Stage interaction survived correction; the analysis therefore did not provide statistically significant evidence of a group difference in early-to-late activity change, and the putaminal group difference remained stable across learning stages.

These results were robust to the prespecified ROI-coverage threshold and were largely unchanged under the stricter coverage sensitivity analysis. Complete ROI statistics and sensitivity analyses are reported in **Tables S7 and S9**.

#### Brain–behavior and connectivity analyses

Five preregistered confirmatory analyses examined associations between performance convergence and learning-related activation, lesion-load moderation, and bilateral M1 functional connectivity. Four analyses yielded no significant findings after voxel-wise FWE correction. The remaining analysis identified a single-voxel lesion-load interaction in one gPPI model that was not reproduced across homologous seed definitions and was therefore considered exploratory. Secondary sensitivity analyses did not alter these conclusions. Results of all preregistered confirmatory analyses are summarized in **Table S8**.

Overall, lesion-aware task-fMRI successfully characterized distributed motor-learning networks, revealing stage-dependent modulation across selected visuomotor regions together with reduced bilateral putaminal activity after stroke. The present sample, however, did not provide reproducible evidence that individual differences in performance convergence were associated with learning-related regional activation or M1–M1 functional connectivity.

## Discussion

This study demonstrates that a sequence-specific reduction in interlimb accuracy asymmetry provides a temporally resolved behavioral phenotype of preserved motor sequence learning in chronic stroke and characterizes its neural context using lesion-aware task-fMRI. Three principal findings emerged. First, stroke survivors demonstrated a progressive, sequence-specific reduction in interlimb accuracy asymmetry, indicating that paretic-hand performance increasingly approached that of the less-affected hand despite persistent motor impairment. Second, lesion-aware neuroimaging revealed preserved stage-dependent modulation within key nodes of the visuomotor sequence-learning network together with reduced bilateral putaminal activity after stroke. Third, no reproducible association was observed between individual differences in this behavioral phenotype and learning-related activation or functional connectivity at the present sample size. Sensitivity analyses supported this interpretation (**Tables S8–S10**).

### Performance convergence during sequence learning

The principal behavioral finding was that stroke participants exhibited greater performance convergence during sequence than random practice. Although interlimb accuracy asymmetry decreased under both conditions, the reduction was significantly greater during sequence learning, indicating a sequence-specific component beyond condition-general practice or session effects.

These findings extend evidence that important components of motor skill and sequence learning remain available after stroke, including in the chronic phase, although their expression varies with task demands, impairment, cognition, and lesion characteristics^5–7,49,50^. Previous work has demonstrated retained sequence-learning capacity despite persistent motor impairment, while studies in neurologically healthy adults show that both behavior and neural activity evolve across separable learning stages^8,9,51^. The present results extend this work by showing that temporally resolved behavioral trajectories reveal learning-related change not captured by conventional endpoint measures. Whereas overall accuracy demonstrated a group-dependent sequence benefit that remained stable across practice, interlimb accuracy asymmetry captured the sequence-specific evolution of performance during learning.

Conventional measures such as average accuracy or overall improvement provide limited insight into how the two hands differentially benefit from learned sequence structure. The greater reduction in interlimb accuracy asymmetry during SEQ than RND indicates that the paretic hand increasingly benefited from predictive sequence knowledge, bringing its accuracy closer to that of the less-affected hand. This sequence-specific performance convergence therefore captures a dynamic aspect of sequence learning beyond endpoint performance and may partly reflect reduced planning disadvantage of the paretic hand during sequential performance^15,16,52^.

Because it can be quantified within a single session, performance convergence provides a practical behavioral phenotype for mechanistic and interventional studies. Improved anticipation of upcoming actions may increase confidence in successful paretic-hand execution, potentially influencing arm-use decisions that are known to depend on anticipated success, effort, habitual nonuse, and cognitive demands^53–58^. Whether performance convergence predicts subsequent hand selection, retention, treatment responsiveness, or real-world arm use remains to be established.

### Neural activity during motor sequence learning

The lesion-aware imaging framework detected robust task-related activation in both stroke and control participants despite substantial anatomical heterogeneity. This finding demonstrates that distributed task-related activity can be characterized in chronic stroke when lesion effects on spatial normalization, voxelwise coverage, and statistical estimability are explicitly addressed.

Whole-brain analyses confirmed expected activation within visual, sensorimotor, and premotor regions during task performance. Although no learning-stage contrast survived whole-brain correction, a priori ROI analyses identified early-to-late decreases in bilateral cerebellar lobule VI, left ventral premotor cortex, bilateral caudate, and right putamen. These changes are broadly consistent with stage-dependent reorganization of premotor, cerebellar, and corticostriatal systems as participants increasingly use learned sequence structure to plan and execute the task. Because motor-sequence learning produces heterogeneous neural responses, the direction of BOLD change alone cannot distinguish improved prediction, reduced error processing, or greater efficiency^9–11,51,59^.

The cerebellar findings are compatible with a changing contribution of internal-model formation, prediction, and online error correction as sequence structure becomes more familiar, whereas the premotor and striatal findings are compatible with changes in advance planning, sequence organization, and corticostriatal action control^10,12,15,52,60^. However, these interpretations remain hypothetical because the present contrasts did not isolate prediction, planning, or error-related computations.

Stroke participants also demonstrated lower bilateral putaminal activity independent of learning stage. Because no Group × Stage interaction was observed, the group and stage effects should be interpreted as distinct findings rather than evidence that stroke altered the temporal trajectory of putaminal activity. The absence of a detectable interaction was consistent with broadly similar early-to-late modulation across groups but does not establish equivalent neural dynamics, and the study was not designed to formally demonstrate such equivalence. Lower putaminal activity may reflect altered corticostriatal recruitment after stroke^5,7,11,20^, but the present data cannot distinguish reduced neural engagement from lesion-related disconnection, compensatory redistribution, neurovascular differences, or task strategy.

### Brain–behavior relationships

The preregistered brain–behavior analyses did not identify reproducible associations between performance convergence and learning-related activation or M1–M1 functional connectivity. One isolated connectivity finding did not replicate across homologous seed definitions and was therefore treated as exploratory.

The analyses readily detected expected task-related activity and regional differences, whereas brain–behavior relationships are particularly sensitive to measurement error, anatomical heterogeneity, and limited sample size. Accordingly, group-average activation does not imply stable between-person brain–behavior associations, particularly in conventional task-fMRI samples^61–64^.

The consistent null pattern across activation, connectivity, and sensitivity analyses therefore provides a study-specific benchmark for the precision achievable with the present design rather than evidence that performance convergence lacks a neural basis. Future studies may improve sensitivity through larger samples, repeated behavioral assessment, greater within-participant imaging data, and individualized anatomical and functional mapping, particularly in stroke where lesion topology increases spatial variability^65,66^.

### Implications for rehabilitation neuroscience

Several limitations constrain interpretation. First, this was a single-site pilot study and was not powered for definitive whole-brain brain–behavior inference. Second, heterogeneous lesion anatomy reduced effective voxel coverage and prevented reliable normalization of one participant’s functional data despite lesion-aware processing. Third, the study measured within-session change at one visit and therefore cannot determine whether performance convergence predicts subsequent retention, longer-term skill consolidation, functional recovery, or spontaneous paretic-arm use^1,7^. Fourth, the selected ROIs and connectivity seeds represented a literature-informed but necessarily restricted set of candidate regions and networks.

Despite these limitations, the study makes complementary conceptual, neurobiological, and methodological contributions. It identifies performance convergence as a temporally resolved behavioral phenotype of preserved motor learning, demonstrates preserved stage-dependent modulation across cerebellar, premotor, and striatal learning networks despite reduced putaminal recruitment after stroke, and establishes lesion-aware task-fMRI as a practical framework for studying these processes in heterogeneous chronic-stroke cohorts.

Rather than simply asking whether performance improves, performance convergence characterizes how the behavioral benefit of predictive sequence knowledge emerges during practice. Future studies should determine whether it generalizes across tasks and sessions, predicts retention^7^ and spontaneous paretic-arm selection^53–58^, and identifies individuals most likely to benefit from rehabilitation or adjunctive neuromodulation^7^.

## Conclusions

Chronic stroke survivors demonstrated sequence-specific performance convergence, expressed as progressive reduction in interlimb accuracy asymmetry during motor sequence learning. Lesion-aware task-fMRI identified robust task-related activation, preserved stage-dependent modulation within distributed visuomotor learning networks, and reduced bilateral putaminal recruitment after stroke, but did not resolve reproducible neural correlates of individual differences in performance convergence at the present sample size. These findings establish performance convergence as a temporally resolved behavioral phenotype beyond conventional endpoint measures and provide a methodological foundation for future studies linking individualized motor learning trajectories to the neural mechanisms that shape paretic-arm use after stroke.

## Data and code availability statement

The complete deidentified dataset — raw BIDS data, lesion masks, and the derivative datasets needed to reproduce the reported analyses — together with all analysis code, will be made publicly available upon publication. The dataset will be released on OpenNeuro (accession ds008676), the analysis code on GitHub, and an archived version deposited on Zenodo with a citable DOI. Materials are available to editors and reviewers during peer review on request.

## Author contributions

Author contributions are reported using the CRediT taxonomy: Conceptualization: K.-F.H., C.F. Methodology: K.-F.H., P.F., P.A.M., C.F., F.K., S.T.S., V.R. Software: K.-F.H., P.A.M., C.F. Validation: K.-F.H., P.F., P.A.M., C.F., F.K., S.H. Formal analysis: K.-F.H., C.F. Investigation: K.-F.H., P.F., P.A.M., F.K., S.H. Resources: K.-F.H., C.F. Data curation: K.-F.H., P.F., P.A.M., C.F., F.K., S.H. Writing – original draft: K.-F.H. Writing – review & editing: K.-F.H., P.F., P.A.M., C.F., F.K., S.H., S.T.S., V.R. Visualization: K.-F.H., C.F., S.T.S. Supervision: K.-F.H., P.F., S.T.S., V.R. Project administration: K.-F.H., P.F., P.A.M., S.H. Funding acquisition: K.-F.H. All authors read and approved the final manuscript.

## Acknowledgements

The authors thank Valerie Salisbury for assistance with participant testing and data acquisition, and Whitney Washington for assistance with lesion-mask delineation. We also sincerely thank all study participants for their time, dedication, and invaluable contribution to this research

