## Supplementary Material for "Sequence-Specific Reduction of Interlimb Accuracy Asymmetry Reveals Preserved Motor Learning Dynamics in Chronic Stroke: Insights from Lesion-Aware fMRI"

### I. SUPPLEMENTARY METHODS

#### Participant flow (Figure S1)

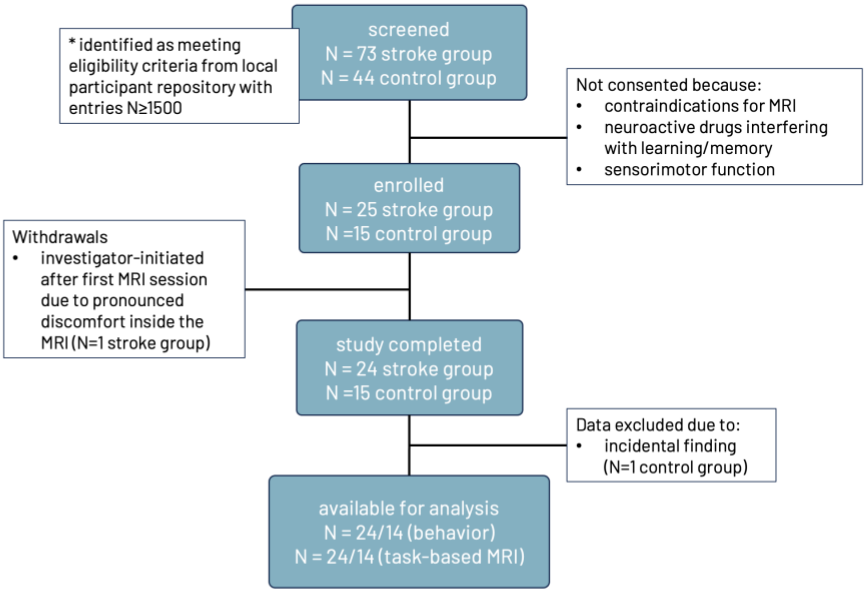

***Figure S1. Participant flow diagram*** *(CONSORT-style). Of N = 73 stroke and N = 44 control individuals identified as meeting eligibility criteria in a local participant repository (≥ 1,500 entries) and screened, N = 25 stroke and N = 15 control participants were enrolled; non-consented individuals were excluded for MRI contraindications, use of neuroactive drugs interfering with learning/memory, or insufficient sensorimotor function. One stroke participant withdrew (investigator-initiated, after the first MRI session, due to pronounced scanner discomfort), yielding N = 24 stroke and N = 15 control participants who completed the study. One control participant was subsequently excluded due to an incidental MRI finding, yielding the final analysis samples of N = 24 stroke and N = 14 control participants for both behavioral and task-based MRI analyses.*

#### Eligibility Screening

Screening was conducted in two stages. First, a standardized telephone screening assessed preliminary eligibility, including stroke history, upper-extremity motor function, cognitive status, and major contraindications to participation. Potentially eligible individuals then completed an in-person screening visit to confirm eligibility regarding the ability to perform the bimanual force-tracking (BFT) task, and routine institutional MRI safety screening.

#### Specific Sensorimotor Eligibility Screening

Maximum voluntary contraction and repeated contraction–relaxation performance were assessed using a digital dynamometer with visual force feedback, employing the same force transducers as used in the experimental learning paradigm. A supplemental visuomotor force-adjustment task was administered under standardized conditions, with participants seated comfortably and forearms supported while visual feedback displayed a target force corresponding to 5% of each participant's maximum voluntary contraction for the left and right hand, alongside real-time feedback of exerted force. Participants completed 20 trials per effector condition for left hand, right hand, and bilateral performance (60 trials total), presented at a variable pacing frequency (0.17–0.25 Hz) with self-paced breaks between blocks (total duration ~6–10 minutes). Decisive eligibility criterion was ≥80% valid trials in each effector condition. After established eligibility, participants signed informed consent including full study procedures.

#### Medication Classification

Self-reported medication data were grouped according to central neuroactive properties and their potential influence on (i) learning/plasticity mechanisms and (ii) fMRI-BOLD signal modulation, based on established pharmacological classes (e.g., GABAergic sedatives, monoaminergic agents, antiseizure medications, stimulants, anticholinergic agents, opioids, gabapentinoids) and physiological modulators of neurovascular coupling (e.g., antihypertensive agents). For each participant, subject-level summary indicators were derived, including binary flags for high neuroactive exposure, high BOLD confound risk, and acute state confound (sedating or stimulant agents), as well as composite burden indices (e.g., sedative load, stimulant load).

#### MRI data preprocessing

Functional and anatomical MRI data were preprocessed using fMRIPrep 25.1.4 (RRID:SCR_016216)[1]. Anatomical preprocessing included intensity non-uniformity correction (N4BiasFieldCorrection)[2], distributed with ANTs 2.6.2 (RRID:SCR_004757)[3], skull-stripping, brain-tissue segmentation into cerebrospinal fluid, white matter, and gray matter (FSL FAST)[4], and cortical surface reconstruction. Rather than reconstructing cortical surfaces within fMRIPrep, FreeSurfer surfaces were generated externally with FreeSurfer 7.4.1 (RRID:SCR_001847)[5] using a standard recon-all on each participant's T1-weighted image and supplied to fMRIPrep as precomputed input. All stroke reconstructions were visually inspected and manually edited to correct reconstruction failures, particularly in and around the lesion, before use; performing surface reconstruction externally is what enabled this manual quality control, ensuring accurate cortical surfaces in structurally heterogeneous brains. Volumetric spatial normalization to MNI152NLin2009cAsym[6,7] was performed via nonlinear registration (antsRegistration, ANTs 2.6.2)[3].

Functional preprocessing included estimation of a BOLD reference volume and head-motion correction (mcflirt)[8], fieldmap-based susceptibility-distortion correction, and boundary-based co-registration to the T1w reference (bbregister)[9]. Confound time series were calculated for framewise displacement[8,10], DVARS, three region-wise global signals (CSF, white matter, whole brain), and anatomical/temporal CompCor components[11], each subsequently expanded with temporal derivatives and quadratic terms[12]; volumes exceeding a framewise-displacement threshold of 0.5 mm or a standardized DVARS threshold of 1.5 were flagged as motion outliers. An additional nuisance-regressor set was derived from a thin band of voxels ("crown") around the brain edge[13]. Preprocessed BOLD runs were resampled into MNI152NLin2009cAsym space (2-mm isotropic resolution) in a single interpolation step combining the head-motion, susceptibility-distortion, and normalization transforms.

#### Lesion Segmentation and Lesion-Aware Analysis Masking

To account for structural brain lesions in the stroke cohort, a complementary workflow was applied alongside the standard fMRIPrep pipeline described above. Stroke lesions were manually segmented on each participant's high-resolution T1-weighted structural scan using ITK-SNAP and reviewed by a board-certified neuroradiologist. Lesion masks were binarized with FSL (fslmaths)[14] and resampled onto the native T1 grid with nearest-neighbor interpolation using ANTs (antsApplyTransforms)[3], ensuring voxel-wise alignment to the T1w image used in fMRIPrep's anatomical workflow.

Spatial normalization was not modified to account for the lesion. The subject-specific T1w to MNI152NLin2009cAsym transform estimated by fMRIPrep was retained and reused to resample the preprocessed BOLD series, the T1w brain mask, and the lesion mask into template space (antsApplyTransforms; linear interpolation for the BOLD series, nearest-neighbor for the masks), yielding functional data and lesion masks that were spatially congruent across subjects. Because the same fMRIPrep transform was applied to every participant, spatial normalization was identical in construction across the stroke and control cohorts.

Lesion information was instead incorporated explicitly at the masking, coverage, and analysis stages, i.e., the sense in which the pipeline is lesion aware. For each stroke participant, an explicit first-level analysis mask was created by subtracting the normalized lesion mask (in MNI space) from the normalized brain mask; this mask was supplied during first-level general linear model estimation (see First-Level GLM and Contrast Definitions, below) so that lesioned voxels were excluded from statistical modeling. Lesion masks in MNI space were stacked to compute overlap and proportion maps, which were visually inspected to verify lesion distributions and to guide construction of the group-level coverage masks (Supplementary Results, Section S6). The same normalized lesion masks were used to derive the whole-brain and M1 lesion-load indices (Lesion-load quantification, below).

#### First-Level GLM and Contrast Definitions

Preprocessed task-fMRI data were imported from the fMRIPrep derivatives into a dedicated SPM25 workflow. For each participant, MNI152NLin2009cAsym-normalized BOLD runs (2-mm isotropic resolution), corresponding confound time series, task events files, and subject-specific brain masks were copied into an SPM analysis tree without modification of the original preprocessing outputs; in stroke participants, spatial normalization had been performed upstream by fMRIPrep, and the lesion-excluded explicit analysis mask described above was applied during model estimation. Task events were converted to SPM multiple-conditions files, and nuisance regressors were extracted from the fMRIPrep confounds tables into SPM-compatible text files. Only experimental task runs were included; localizer runs were excluded a priori.

Within each participant, runs were resliced to a common reference grid (first retained run) to ensure identical image geometry across runs and were subsequently smoothed using an isotropic Gaussian kernel (6 mm full width at half-maximum). To constrain model estimation to valid functional tissue, a subject-specific explicit mask was constructed by combining the fMRIPrep-derived analysis mask (lesion-excluded, for stroke participants) with a data-driven coverage criterion: voxels were retained if they exhibited valid signal in at least 80% of sampled volumes across runs and exceeded a conservative intensity threshold based on the run-averaged signal. Runs containing volumes with no overlap with the subject mask were excluded, and the explicit mask was recomputed from the remaining valid runs.

First-level statistical analyses were performed in SPM25 using a general linear model with one session per run. The design matrix included two task conditions, sequential learning (SEQ) and random control (RND), modeled using the canonical hemodynamic response function (HRF) with temporal derivatives. Low-frequency signal drifts were removed using a discrete cosine transform basis set with a high-pass filter cutoff of 128 s.

Nuisance effects were modeled following standard SPM conventions. Head motion was accounted for by including the six rigid-body realignment parameters (three translations and three rotations) together with their temporal derivatives, yielding 12 motion regressors per run. Volumes identified as motion or intensity outliers in the fMRIPrep confounds were modeled using separate single-volume ("spike") regressors. All nuisance regressors were entered as run-specific covariates of no interest. Serial correlations in the BOLD time series were modeled using a first-order autoregressive [AR(1)] process with restricted maximum likelihood estimation. Model estimation was restricted to the subject-specific explicit mask described above; in cases where estimation failed due to insufficient estimable voxels, the model was re-estimated assuming independent errors.

Contrast estimates were computed from the final model after any run exclusions, to ensure valid weighting of retained data. First-level contrasts included the main effects of SEQ and RND across runs, their difference (SEQ−RND), and time-related contrasts capturing change between early and late task periods, as well as their combinations (e.g., the early-to-late change in the sequence-specific SEQ−RND effect). Each contrast was estimated in both positive and negative directions for second-level inference. Subject-level contrast images were subsequently entered into the second-level random-effects analyses described in the main manuscript and above (Supplementary Results, Section S6).

#### Behavioral preprocessing

Grip force was recorded at 1000 Hz and converted to relative force (% of target force). Trials with missing or invalid force data were excluded. The preregistered primary analyses (PRIMARY) reported in the main manuscript (Models A–C and the hand-specific accuracy-lag models D1–D3) use two trial-level metrics computed across the full trial trajectory: accuracy, the root-mean-square (RMS) deviation from target force across the trial (rmsErrorTrial; higher values indicate worse, less accurate tracking), and reaction time (RT), the latency to reach the target force level.

#### Primary performance-convergence model

For each trial, an accuracy lag was defined as the between-hand difference in RMS force-tracking error (affected/paretic hand minus less-affected hand; an analogous non-dominant-hand assignment was used in controls; higher values indicate worse, less accurate tracking). Three nested mixed-effects models were fit in R using the same baseline-normalization and random-effects-selection conventions described above for the sequence-benefit models. In all models, Run was modeled as a continuous, mean-centered predictor (rather than as a categorical factor); trial-level observations were nested within participant, and each model included a by-subject random intercept together with, where supported by Akaike Information Criterion and likelihood-ratio comparison, a by-subject random slope for Run (the selected structure refit by restricted maximum likelihood).

- Model D1 regressed the accuracy lag on Condition (SEQ, RND) and Group, with Group × Condition as the primary test term.
- Model D2 extended Model D1 by adding Run and its interactions (Condition × Run, Group × Condition × Run); the three-way term is the pre-specified PRIMARY behavioral test (Table S3).
- Model D3 restricted the Group × Run interaction test to RND trials alone (Table S3).

A per-subject linear performance-convergence slope across runs (lag_catchup_slope_acc) was derived from the SEQ-condition model coefficients and winsorized at the sample median ± 3 median absolute deviations to limit the influence of extreme values; both the winsorized and raw (non-winsorized) slopes were retained for the corresponding PRIMARY/SENSITIVITY fMRI covariate models described in the main manuscript.

#### Secondary Behavioral Models (Models A–C)

Overall sequence benefit and its evolution across runs were characterized using trial-level accuracy and RT models (see Behavioral preprocessing, above, for outcome definitions) that include hand role (affected, less-affected) as a fixed-effect covariate rather than as the outcome itself complementing the hand-specific accuracy-lag models (D1–D3, below), which take the between-hand difference itself as the outcome. All models include, subject as a random effect; candidate random-effects structures (intercept only; intercept plus correlated or uncorrelated random slope for Run) were compared by Akaike Information Criterion and likelihood-ratio test, with the selected structure refit by restricted maximum likelihood, following the same conventions used for Models D1–D3. Full fixed-effect statistics for all three models are reported in Table S5.

- Model A regressed accuracy and RT on Condition (SEQ, RND), Group (Stroke, Control), Hand role, Run (mean-centered, main effect only), Block order, and the corresponding baseline covariate, with Group × Condition as the model's primary test term.
- Model B extended Model A by adding Condition × Run and Group × Condition × Run interaction terms, with the three-way term as the model's primary test (cf. Model D2, below).

- Model C regressed RND-only accuracy and RT on Group, Timepoint (pre-/post-task localizer), Group × Timepoint, and the corresponding baseline covariate, with a random intercept for subject; unlike Models A and B, Model C uses the localizer runs rather than the SEQ/RND task runs.

Model diagnostics (residual behavior, collinearity, random-effects singularity) and Type III analyses of variance for fixed effects followed standard practice for all models above.

#### Lesion-load quantification (extended methods)

Whole-brain lesion-load fraction was computed per stroke participant as the proportion of the canonical brain-tissue mask (see main manuscript) classified as non-brain within that participant's first-level analysis mask. A region-restricted variant was computed identically within a bilateral M1 hand-knob mask.

To reduce non-lesion-related variability arising from imperfect tissue coverage, lesion-load estimates were normalized using the median value observed in controls. The corrected whole-brain metric was used as the default lesion-load index in the second-level moderation model described in the main manuscript; the raw (uncorrected) metric was retained as a SENSITIVITY comparison.

One participant (excluded from all imaging analyses; see main manuscript Participants) was identified during quality control to have a genuine but small, previously unsegmented lesion; a second participant's lesion mask could not be reliably registered to template space (a large majority of canonical brain tissue flagged as non-brain, implausible for the documented lesion and inconsistent with independent structural imaging) and was excluded from the stroke imaging cohort entirely rather than assigned a lesion-load value.

The bilateral M1 hand-knob-restricted lesion fraction was zero for every stroke participant, consistent with this cohort's lesions being predominantly non-cortical/subcortical and sparing the hand-knob region rather than reflecting a measurement error, consistent with this region successfully showed significant small-volume-corrected activation elsewhere in these analyses.

#### Functional connectivity (gPPI) — extended methods

*Seed regions of interest.* The PRIMARY seed pair was bilateral primary motor cortex (M1 hand-knob; 10-mm-radius sphere, MNI coordinates as used for the corresponding ROI analysis in the main manuscript), defined separately as ipsilesional and contralesional relative to each stroke participant's affected hand. Hemisphere assignment used the clinically coded ipsilesional-hemisphere field in the study's participant data dictionary; one stroke participant with documented multifocal/bilateral lesions had no single defined ipsilesional hemisphere and was excluded from all lateralized-seed (M1, putamen, superior-parietal) analyses while remaining in the cohort for midline seeds and all non-connectivity analyses. An analogous non-dominant-hand convention was used to assign pseudo-affected laterality in controls.

Four additional, secondary/exploratory seed regions were defined: pre-supplementary motor area (midline), bilateral putamen (motor-learning coordinate), bilateral superior parietal lobule/intraparietal sulcus, and dorsal anterior cingulate/cingulate motor area (midline), at the same coordinates used for the corresponding ROI analyses in the main manuscript (10-mm radius).

VOI extraction. For each seed, subject, and run, the first eigenvariate of the BOLD signal was extracted using SPM25's non-interactive volume-of-interest batch job (spm.util.voi) with a fixed geometric (spherical) region definition, following estimation of an effects-of-interest F-contrast for each subject's first-level model. Voxel-coverage diagnostics were retained per seed/session; secondary lateralized seeds occasionally had reduced or zero usable coverage in participants whose lesion directly overlapped that seed's sphere, which is reported as a per-seed data limitation rather than excluded post hoc from the affected analyses.

PPI first-level model. Condition-specific psychophysiological interaction terms (SEQ, RND) were computed for each seed and session using the standard deconvolution approach (spm_peb_ppi) and entered into an augmented first-level general linear model alongside the seed's own BOLD eigenvariate (covariate of no interest), the original task regressors, motion/outlier regressors, and session constants.

#### Analysis pipeline and full list of pre-specified vs. exploratory tests

Table S1 lists every formally pre-specified PRIMARY test alongside its SENSITIVITY/SPECIFICITY counterparts and the broader secondary/exploratory analyses referenced in the main manuscript Statistical framework section.

##### Table S1: Analysis pipeline of pre-specified vs. exploratory tests

| *Family* | *PRIMARY test* | *SENSITIVITY/ SPECIFICITY variants* | *Secondary/ exploratory* |
|---|---|---|---|
| *Activation GLM — bivariate covariate* | SEQ−RND Early-minus-Late, stroke, winsorized covariate | Raw covariate (stroke); winsorized covariate (controls) | SEQ-only and RND-only contrasts |
| *Activation GLM — lesion-load moderation* | Lag × corrected-lesion-load interaction | Raw (uncorrected) lesion-load metric | Main effects of lag and lesion load |
| *ROI / small-volume correction* | — (not a standalone PRIMARY test; applied to PRIMARY contrasts above) | — | M1 hand-knob, SMA, putamen, SPL/IPS, dACC ROIs × all contrasts |
| *gPPI — learning-related change* | SEQ/RND Early-minus-Late, M1–M1 seed pair | — | SMA, putamen, SPL/IPS, dACC seeds |
| *gPPI — behavioral covariate* | SEQ−RND Early-minus-Late × lag, M1–M1 seed pair | — | SEQ-only and RND-only covariate contrasts |
| *gPPI — lesion-load moderation* | Lag × corrected-lesion-load interaction, M1–M1 seed pair | Raw lesion-load metric | Main effects of lag and lesion load |
| *Second-level smoothing* | 6-mm (first-level-smoothed only) | 8-mm additional group-level smoothing (all families) | — |

| | | | |
|---|---|---|---|
| *Early/Late time window* | 1-run window (first vs. last run) | 2-run-block window | — |

#### Table S2: ROI and functional-connectivity seed coordinates

Anatomical ROIs (primary motor cortex, superior parietal lobule, caudate, putamen, lateral occipital cortex) were drawn from the Harvard–Oxford atlases as described in the main manuscript. Coordinate-based spherical ROIs and connectivity seeds are listed below; the M1 hand-knob, SMA, putamen, SPL/IPS, and dACC entries double as both the ROI-analysis spheres (6 mm, main manuscript) and the gPPI connectivity seeds (10 mm, main manuscript and Supplementary Methods).

| ***Region*** | ***MNI coordinate(s)*** | ***Radius*** | ***Role*** | ***Source*** |
|---|---|---|---|---|
| *Pre-supplementary motor area (pre-SMA)* | [0, 17, 55] | 6 mm | ROI (stage-dependent profile) | Mayka et al. 2006[15] |
| *Dorsal premotor cortex (PMd)* | [±32, –12, 60] | 6 mm | ROI (stage-dependent profile) | Hardwick et al. 2013[16] |
| *Ventral premotor cortex (PMv)* | [±53, 12, 25] | 6 mm | ROI (stage-dependent profile) | Mayka et al. 2006[15] |
| *Cerebellar lobule VI* | [±32, –52, –28] | 6 mm | ROI (stage-dependent profile) | Hardwick et al. 2013[16] |
| *M1 hand-knob (ipsi-/contralesional)* | [±38,–24,50] (approx., mirrored by hemisphere) | 10 mm | PRIMARY gPPI seed; ROI/SVC | Mayka et al. 2006[15]; mirrored per participant's ipsilesional hemisphere |
| *SMA proper* | [0,-6,56] (midline) | 10 mm | Secondary gPPI seed; SVC ROI | Hardwick et al. 2013[16] |
| *Striatum — putamen (motor-learning peak)* | [±28,4,4] | 10 mm | Secondary gPPI seed; SVC ROI | Hardwick et al. 2013[16] |
| *Parietal SPL/IPS* | [±30,-52,50] | 10 mm | Secondary gPPI seed; SVC ROI | Hardwick et al. 2013[16] |
| *Cingulate — dACC/CMA* | [0,6,38] (midline) | 12 mm | Secondary gPPI seed; SVC ROI | Mayka et al. 2006[15] |

### II. SUPPLEMENTARY RESULTS

#### Behavior

##### S1. Primary performance-convergence model (full statistics)

All models use accuracy lag (affected minus less-affected hand, RND-baseline-corrected) as the outcome; n = 24 stroke, 14 control (behavioral sample). D1 tests the overall inter-hand asymmetry and its relation to condition; D2 (PRIMARY) adds Run to test whether the asymmetry narrows across the session and whether it does so differently by group; D3 restricts to RND-only localizer runs (pre/post) as a condition-general drift control.

#### Table S3: Full statistics — hand-specific accuracy-lag models (D1–D3)

| MODEL | EFFECT | X²(DF) | P |
|---|---|---|---|
| D1 — SEQUENCE BENEFIT (LAG) | Group | 0.46(1) | .499 |
| D1 — SEQUENCE BENEFIT (LAG) | Condition | 0.04(1) | .844 |
| D1 — SEQUENCE BENEFIT (LAG) | Run | 0.97(1) | .325 |
| D1 — SEQUENCE BENEFIT (LAG) | Block order | 1.20(1) | .273 |
| D1 — SEQUENCE BENEFIT (LAG) | Group × Condition | 4.36(1) | .037 |
| D2 — LEARNING ACROSS EXPOSURE (LAG), PRIMARY | Group | 0.02(1) | .898 |
| D2 — LEARNING ACROSS EXPOSURE (LAG), PRIMARY | Condition | 0.04(1) | .837 |
| D2 — LEARNING ACROSS EXPOSURE (LAG), PRIMARY | Run | 0.00(1) | .977 |
| D2 — LEARNING ACROSS EXPOSURE (LAG), PRIMARY | Block order | 16.20(1) | <.001 |
| D2 — LEARNING ACROSS EXPOSURE (LAG), PRIMARY | Group × Condition | 4.74(1) | .029 |
| D2 — LEARNING ACROSS EXPOSURE (LAG), PRIMARY | Group × Run | 0.02(1) | .875 |
| D2 — LEARNING ACROSS EXPOSURE (LAG), PRIMARY | Condition × Run | 0.00(1) | .999 |
| D2 — LEARNING ACROSS EXPOSURE (LAG), PRIMARY | Group × Condition × Run | 5.76(1) | .016 |
| D3 — LOCALIZER DRIFT CONTROL (LAG) | Group | 3.70(1) | .054 |
| D3 — LOCALIZER DRIFT CONTROL (LAG) | Timepoint | 0.16(1) | .691 |
| D3 — LOCALIZER DRIFT CONTROL (LAG) | Group × Timepoint | 7.47(1) | .006 |

##### S2. Secondary behavioral models (full statistics)

#### Table S4: Full behavioral model statistics (Models A–C)

Full fixed-effect statistics for the secondary sequence-benefit and drift models summarized in the main manuscript Results (Behavior): trial-level accuracy and RT, modeled with hand role (affected, less-affected) as a fixed-effect covariate rather than as the outcome itself (see Methods).

Model D1–D3 (PRIMARY hand-specific accuracy-lag models) full statistics are reported separately in Table S3 (Section S1).

| MODEL | OUTCOME | EFFECT | X²(DF) | P | R² (COND./MARG.) |
|---|---|---|---|---|---|
| A — SEQUENCE BENEFIT | Accuracy | Hand role | 96.56(1) | <.001 | 0.15 / 0.02 |
| A — SEQUENCE BENEFIT | Accuracy | Condition | 0.01(1) | .936 | |
| A — SEQUENCE BENEFIT | Accuracy | Group | 1.68(1) | .195 | |
| A — SEQUENCE BENEFIT | Accuracy | Group × Condition | 12.99(1) | <.001 | |
| A — SEQUENCE BENEFIT | Accuracy | Run | 8.01(1) | .005 | |
| A — SEQUENCE BENEFIT | Accuracy | Block order | 9.44(1) | .002 | |
| A — SEQUENCE BENEFIT | RT | Hand role | 45.65(1) | <.001 | 0.29 / 0.21 |
| A — SEQUENCE BENEFIT | RT | Condition | 0.14(1) | .712 | |
| A — SEQUENCE BENEFIT | RT | Group | 0.12(1) | .729 | |
| A — SEQUENCE BENEFIT | RT | Group × Condition | 2.70(1) | .100 | |
| A — SEQUENCE BENEFIT | RT | Run | 8.43(1) | .004 | |
| A — SEQUENCE BENEFIT | RT | Block order | 10.91(1) | <.001 | |
| B — LEARNING ACROSS EXPOSURE | Accuracy | Hand role | 99.00(1) | <.001 | 0.19 / 0.02 |
| B — LEARNING ACROSS EXPOSURE | Accuracy | Condition | 0.01(1) | .934 | |
| B — LEARNING ACROSS EXPOSURE | Accuracy | Group × Condition | 13.52(1) | <.001 | |
| B — LEARNING | Accuracy | Block order | 35.61(1) | <.001 | |

| | | | | | |
|---|---|---|---|---|---|
| **ACROSS EXPOSURE** | | | | | |
| **B — LEARNING ACROSS EXPOSURE** | Accuracy | Group × Condition × Run | 0.08(1) | .780 | |
| **B — LEARNING ACROSS EXPOSURE** | RT | Hand role | 39.80(1) | <.001 | — / 0.24 |
| **B — LEARNING ACROSS EXPOSURE** | RT | Condition | 0.12(1) | .734 | |
| **B — LEARNING ACROSS EXPOSURE** | RT | Group × Condition | 2.35(1) | .126 | |
| **B — LEARNING ACROSS EXPOSURE** | RT | Block order | 5.99(1) | .014 | |
| **B — LEARNING ACROSS EXPOSURE** | RT | Group × Condition × Run | 1.18(1) | .278 | |
| **C — SESSION DRIFT** | Accuracy | Hand role | 6.93(1) | .008 | 0.22 / 0.18 |
| **C — SESSION DRIFT** | Accuracy | Group | 5.94(1) | .015 | |
| **C — SESSION DRIFT** | Accuracy | Timepoint | 9.84(1) | .002 | |
| **C — SESSION DRIFT** | Accuracy | Group × Timepoint | 2.47(1) | .116 | |
| **C — SESSION DRIFT** | RT | Hand role | 0.62(1) | .432 | 0.32 / 0.29 |
| **C — SESSION DRIFT** | RT | Group | 0.92(1) | .338 | |
| **C — SESSION DRIFT** | RT | Timepoint | 14.21(1) | <.001 | |
| **C — SESSION DRIFT** | RT | Group × Timepoint | 7.33(1) | .007 | |

#### S3. Follow-up simple-effects and slope contrasts

The omnibus tests in Tables S3–S4 establish that the Group × Condition, Group × Condition × Run, and Group × Timepoint interactions are statistically significant but do not by themselves identify which specific comparison drives each effect. Table S5 reports the corresponding simple-effects and slope follow-up contrasts, computed directly on the fitted models underlying Tables S3–S4 (emtrends for run-slopes; emmeans for simple effects; Holm-corrected within each family of planned comparisons). Given trial-level sample sizes exceeding lmerTest's default denominator-df threshold, these follow-up tests use asymptotic z rather than Satterthwaite t; the two are numerically equivalent at these sample sizes.

For the PRIMARY Group × Condition × Run interaction (Model D2, Table S3), no single Group × Condition cell's run-slope differed significantly from zero, including the stroke/SEQ cell (slope = −1.23, SE = 1.89, z = −0.65, p = .515). The interaction was instead driven by a significant condition-specific slope difference within the stroke group (SEQ vs. RND: estimate = 1.79, SE = 0.45, z = 3.95, p < .001); the direct stroke-SEQ vs. control-SEQ slope contrast did not reach significance (estimate = 1.30, SE = 3.11, z = 0.42, p = 1.000). The reduction in interlimb asymmetry across runs is therefore best characterized as a condition-specific (sequence-practice-selective) effect within the stroke group, rather than an independently significant divergence from the control group's trajectory.

For the Group × Condition interaction at initial exposure (Model D1), the simple stroke-vs-control contrast was not significant within either condition (RND: p = .499; SEQ: p = .222).

For the Group × Timepoint interaction on the RND-localizer accuracy lag (Model D3), stroke participants showed a significant reduction in interlimb asymmetry from pre- to post-task assessment (estimate = 2.95, SE = 0.74, z = 3.98, p < .001), while control participants showed no change (p = .691); the two groups differed marginally at the pre-task assessment (p = .054) and were statistically indistinguishable by the post-task assessment (p = .789).

For the Group × Timepoint interaction on RT in the localizer runs (Model C), control participants slowed significantly from pre- to post-task assessment (estimate = −0.037 s, SE = 0.010, z = −3.77, p < .001), while stroke participants showed no significant change (p = .746).

For the secondary Group × Condition interaction on trial-level accuracy (Model A), the simple stroke-vs-control contrast did not reach significance within either condition (RND: p = .195; SEQ: p = .077).

#### Table S5: Simple-effects and slope follow-up contrasts (Tables S3–S4)

| MODEL | CONTRAST | ESTIMATE | SE | Z | P (HOLM) |
|---|---|---|---|---|---|
| **D2 ACCURACY LAG (PRIMARY)** | Slope vs 0: Control, RND | 0.07 | 2.47 | 0.03 | .977 |
| **D2 ACCURACY LAG (PRIMARY)** | Slope vs 0: Stroke, RND | 0.56 | 1.89 | 0.30 | .767 |
| **D2 ACCURACY LAG (PRIMARY)** | Slope vs 0: Control, SEQ | 0.07 | 2.47 | 0.03 | .977 |
| **D2 ACCURACY LAG (PRIMARY)** | Slope vs 0: Stroke, SEQ | −1.23 | 1.89 | −0.65 | .515 |

| | | | | | |
|---|---|---|---|---|---|
| **D2 ACCURACY LAG (PRIMARY)** | Slope contrast: Control RND − Stroke RND | −0.49 | 3.11 | −0.16 | 1.000 |
| **D2 ACCURACY LAG (PRIMARY)** | Slope contrast: Control RND − Control SEQ | 0.00 | 0.59 | 0.00 | 1.000 |
| **D2 ACCURACY LAG (PRIMARY)** | Slope contrast: Control RND − Stroke SEQ | 1.30 | 3.11 | 0.42 | 1.000 |
| **D2 ACCURACY LAG (PRIMARY)** | Slope contrast: Stroke RND − Control SEQ | 0.49 | 3.11 | 0.16 | 1.000 |
| **D2 ACCURACY LAG (PRIMARY)** | Slope contrast: Stroke RND − Stroke SEQ | 1.79 | 0.45 | 3.95 | <.001 |
| **D2 ACCURACY LAG (PRIMARY)** | Slope contrast: Control SEQ − Stroke SEQ | 1.30 | 3.11 | 0.42 | 1.000 |
| **D1 ACCURACY LAG** | Simple group effect (Control−Stroke), RND | −3.43 | 5.08 | −0.68 | .499 |
| **D1 ACCURACY LAG** | Simple group effect (Control−Stroke), SEQ | −6.21 | 5.08 | −1.22 | .222 |
| **D3 ACCURACY LAG LOCALIZER DRIFT** | Simple timepoint effect (Pre−Post), Control | −0.39 | 0.97 | −0.40 | .691 |
| **D3 ACCURACY LAG LOCALIZER DRIFT** | Simple timepoint effect (Pre−Post), Stroke | 2.95 | 0.74 | 3.98 | <.001 |
| **D3 ACCURACY LAG LOCALIZER DRIFT** | Simple group effect (Control−Stroke), Pre | −2.93 | 1.52 | −1.92 | .054 |
| **D3 ACCURACY LAG LOCALIZER DRIFT** | Simple group effect (Control−Stroke), Post | 0.41 | 1.52 | 0.27 | .789 |
| **MODEL C RT LOCALIZER DRIFT** | Simple timepoint effect (Pre−Post), Control | −0.037 | 0.010 | −3.77 | <.001 |
| **MODEL C RT LOCALIZER DRIFT** | Simple timepoint effect (Pre−Post), Stroke | −0.003 | 0.008 | −0.32 | .746 |

| | | | | | |
|---|---|---|---|---|---|
| **MODEL C RT LOCALIZER DRIFT** | Simple group effect (Control−Stroke), Pre | −0.018 | 0.019 | −0.96 | .338 |
| **MODEL C RT LOCALIZER DRIFT** | Simple group effect (Control−Stroke), Post | 0.016 | 0.019 | 0.85 | .398 |
| **MODEL A ACCURACY (SECONDARY)** | Simple group effect (Control−Stroke), RND | −8.46 | 6.53 | −1.30 | .195 |
| **MODEL A ACCURACY (SECONDARY)** | Simple group effect (Control−Stroke), SEQ | −11.56 | 6.53 | −1.77 | .077 |

All p values Holm-corrected within each family of planned comparisons (D2 run-slope contrasts: 6 comparisons; all other rows: 2 comparisons per model). Estimates are in the original outcome units (accuracy-lag models: %MVC RMS-force-error difference, affected − less-affected hand; RT/RT-lag models: seconds).

#### MRI Quality Control

##### S4. Motion and preprocessing QC

Framewise displacement (FD)[14] was computed by fMRIPrep from the six rigid-body realignment parameters for every task-learn BOLD run, excluding the two runs (run-00, run-07, i.e., the pre-/post localizer runs) already excluded at first-level modeling (Methods). Descriptives below cover all participants with usable fMRIPrep confounds output: 24 stroke participants (144 runs) and 14 control participants (84 runs). Two enrolled participants had no usable confounds output and are not included below: sub-LRN015 (no functional data) and sub-LRN029 (excluded from all analyses on unrelated coverage grounds; see Methods, Participants). This motion-QC sample also includes sub-LRN010, whose data were subsequently excluded from all whole-brain imaging analyses for a separate, unrelated reason (failed spatial normalization driven by multiple large lesions; see Methods): sub-LRN010's subject-mean FD (0.356 mm) sits at the 58th percentile of the N=24 stroke distribution summarized here, i.e. unremarkable relative to the rest of the cohort, consistent with normalization, not motion, having driven that exclusion.

Group-level descriptive statistics are summarized in Table S6. Motion was, as expected for a stroke cohort, somewhat higher in the stroke group (median run-level mean FD = 0.330 mm) than in controls (0.262 mm), with a correspondingly higher proportion of stroke runs exceeding the primary FD threshold (median 1.2% vs. 0.3% of volumes per run); the large majority of runs in both groups fell well within this threshold. fMRIPrep's combined FD/DVARS outlier flags[17,18] were, per Methods, included as single-volume spike regressors at the first level.

#### Table S6: Group-level head-motion quality-control summary

| *Metric* | *Stroke* | *Controls* |
|---|---|---|
| *Mean FD (mm), median [range]* | 0.330 [0.136–0.945] | 0.262 [0.107–0.515] |
| *Mean FD (mm), mean (SD)* | 0.356 (0.147) | 0.264 (0.098) |
| *Max FD (mm), median [range]* | 1.812 [0.390–10.781] | 1.118 [0.361–5.339] |
| *Volumes with FD > 1.0 mm[a] (%), median [range]* | 1.2 [0.0–34.1] | 0.3 [0.0–6.3] |
| *Outlier frames[b] per run, median [range]* | 44 [0–240] | 18 [0–131] |

*[a]Primary threshold = 0.5× resampled voxel size (2 mm) = 1.0 mm. [b]Combined FD/DVARS outlier flags from fMRIPrep's motion_outlier* confound columns, included as single-volume spike regressors at the first level (Methods).*

One stroke participant, sub-LRN007, showed clearly elevated motion relative to the rest of the cohort (subject-mean FD = 0.675 mm; mean 19.9% of volumes/run exceeding 1.0 mm, against a cohort median of 1.2%) and was the largest outlier on every run-level motion metric reported here. Sub-LRN007 was retained in the primary analyses; the spike regression described above mitigates the influence of the specific flagged high-motion frames within this participant's data, but this participant contributes the most motion-affected data in the stroke cohort. A separate, isolated event was noted for sub-LRN021 (a single run reaching a maximum FD of 10.78 mm despite an otherwise-unremarkable subject-mean FD of 0.48 mm), consistent with one brief head movement rather than sustained elevated motion.

#### S5. ROI coverage diagnostics

ROI coverage (proportion of valid, non-lesioned voxels contributing to each subject's ROI-average contrast estimate) was quantified for all 17 a priori ROIs (Supplementary Table S2) across 24 stroke participants and all task-learn contrasts, following the same procedure used to enforce the ≥25% minimum-coverage rule at first pass (Methods).

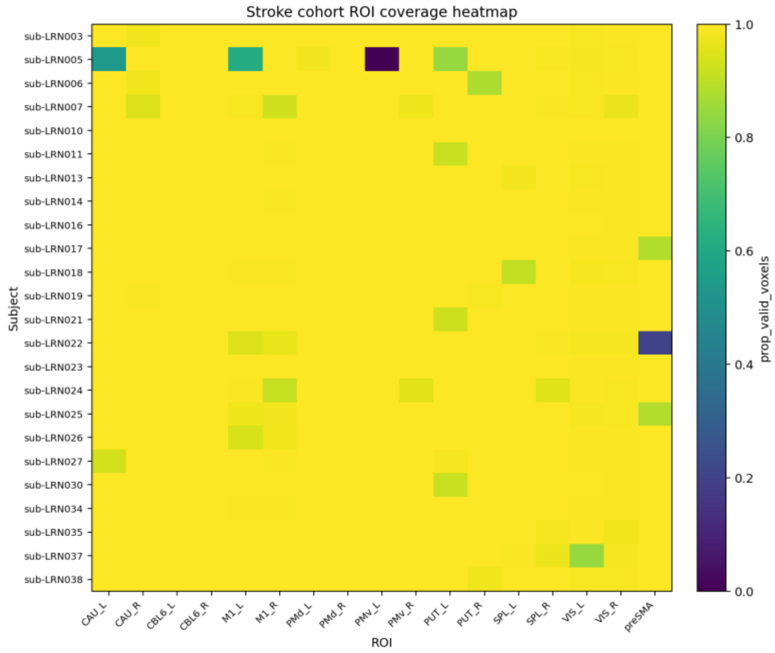

***Figure S5a.*** *Subject × ROI coverage heatmap, stroke cohort (mean proportion of valid voxels per ROI, averaged across contrasts). Two cells stand out as near-complete non-coverage: sub-LRN005 in left ventral premotor cortex (PMv_L) and sub-LRN022 in pre-supplementary motor area (preSMA).*

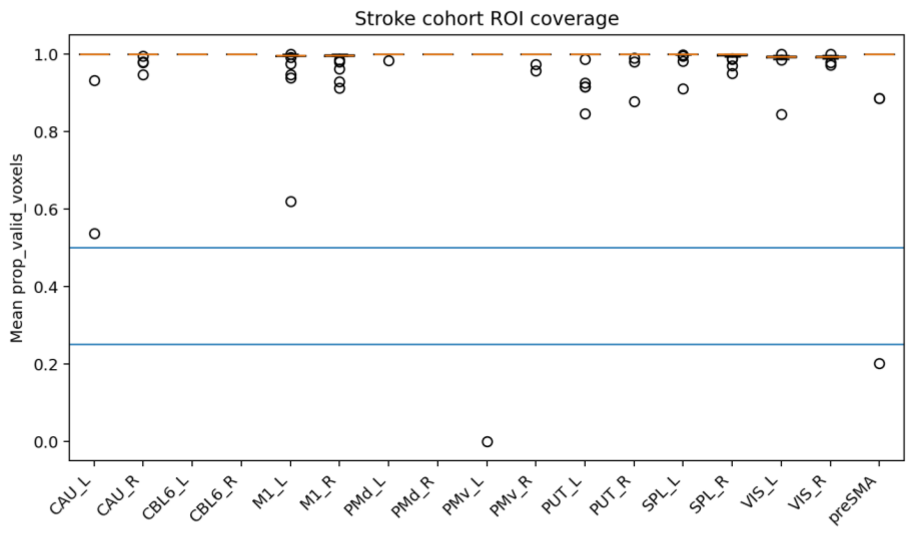

***Figure S5b.*** *Per-ROI coverage distributions, stroke cohort (each point = one subject's mean coverage in that ROI). Horizontal lines mark the 25% and 50% minimum-coverage thresholds (Methods); all values below are the same two cells as Figure S5a.*

Coverage was high overall: median coverage was ≥99% of voxels in 16 of 17 ROIs, and only 2 of 408 subject×ROI cells (0.5%) fell below the 25% threshold — both isolated to single subjects (sub-LRN005 in left PMv, 0% valid voxels; sub-LRN022 in preSMA, 20.2% valid voxels), each affecting all 7 contrasts for that subject×ROI combination and excluded from the corresponding mixed-effects models under the pre-specified rule (Methods). Neither the affected ROIs nor any other ROI showed a broader pattern of degraded coverage across the cohort.

At the same time, per-subject mean coverage (averaged across all 17 ROIs) showed a moderate negative relationship with whole-brain lesion load[14,17] (lesion_frac_wholebrain_corrected; *Spearman* $\rho = -0.48$, $p = .021$, $N = 23$; sub-LRN010 has no imaging-analysis lesion-load estimate due to failed spatial normalization and is not included here, see Methods). This association is not solely an artifact of the single highest-lesion-load subject: excluding sub-LRN005 (whose lesion load is roughly 4× the next-highest subject) it remains directionally consistent and remains significant by a parametric test (Pearson $r = -0.54$, $p = .010$, $N = 22$; Spearman $\rho = -0.41$, $p = .062$). This is the anatomically expected direction (larger lesions remove more voxels from the explicit analysis mask near the lesion) and is consistent with the two isolated low-coverage cases above both occurring in relatively high-lesion-load subjects (sub-LRN005 ranks first and sub-LRN022 ranks third of 23 stroke participants by lesion load).

Taken together, these diagnostics do not support a blanket claim that lesion effects are absent from ROI extraction. Instead, a modest, expected coverage–lesion-load relationship was observed. They do show that this relationship is small in magnitude, concentrated in two identifiable subject×ROI cases rather than distributed across the ROI set, and already handled by the pre-specified ≥25% coverage exclusion rule (with results reported as consistent under the stricter ≥50% threshold;

Methods). No evidence was found that residual lesion-driven coverage loss biases the reported ROI results beyond the two excluded cases already documented here.

#### MRI Results

##### S6. Whole-brain statistics

Whole-brain second-level results are summarized here for the task-positive validation contrasts (SEQ, RND, and their conjunction) and the learning-related contrasts (Early-minus-Late; SEQ−RND at Early, Late, and Early-minus-Late), at the primary analysis level (6-mm first-level smoothing, no additional group-level smoothing) and under an additional 8-mm group-level smoothing arm used as a sensitivity check (Methods).

Task-positive validation contrasts (SEQ, RND, conjunction) showed robust, anatomically expected voxel-wise FWE-corrected ($p<.05$) clusters in both cohorts at the primary smoothing level (Figure 2B–C).

Learning-related contrasts (Early-minus-Late, and SEQ−RND at Early or Late) were null at voxel-wise FWE correction in both cohorts at the primary smoothing level. Under the additional 8-mm group-level smoothing arm, one cluster reached FWE significance: SEQ Early-minus-Late, negative direction (i.e., greater late-stage than early-stage activity), controls only: 1 cluster, 2 voxels, peak $Z=11.21$, MNI [-4.5,27.5,5.5]. This cluster was not present in the same cohort and contrast at the primary 6-mm smoothing level (peak uncorrected $Z=6.64$ at a nearby but distinct location, MNI [-0.5,29.5,9.5]; 0 voxels survived FWE correction), and no corresponding cluster reached significance in stroke at either smoothing level. Given its small size (2 voxels), its dependence on a single non-primary smoothing parameterization, and its absence in the stroke cohort, this cluster is not considered a robust finding.

###### *Group coverage maps and lesion-frequency relationship*

Following the coverage-thresholding procedure described in the main manuscript Methods (≥50% of participants in the stroke group, ≥80% in the control group, intersected with the canonical brain-tissue mask), Figure S6 shows the resulting contrast-specific coverage map for the stroke cohort's SEQ+RND all-runs conjunction contrast, i.e., the same task-positive validation contrast already shown for its activation results in main-text Figure 2B, alongside a lesion-frequency map computed over the same matched N=23 functional-analysis sample. Coverage was high overall: of 242,861 in-mask voxels, only 4.4% fell below the 0.50 stroke coverage threshold. The lesion-frequency map shown is derived from the same FSL/lesionnorm pipeline used for the whole-cohort lesion map in main-text Figure 2A (percentage of participants with lesion involvement per voxel), restricted here to the matched N=23 sample by excluding the one participant present in the N=24 lesion-imaging sample but not in the functional-analysis sample (main manuscript, Participants). Coverage and lesion frequency were only weakly related across space in this comparison using the matched analysis sample (Pearson $r=0.06$ within the matched N=23 sample), suggesting that voxel coverage in this pipeline reflects additional exclusion criteria beyond lesion location alone. Note that the lesion-frequency map shown here (N=23) is deliberately the matched functional-analysis sample, not the N=24 lesion-frequency map in main-text Figure 2A; the two lesion maps are visually similar but not numerically interchangeable. The identical coverage/threshold procedure was applied to every other contrast and to the control cohort (≥80% threshold); the representative contrast shown here is representative of the pattern seen across contrasts.

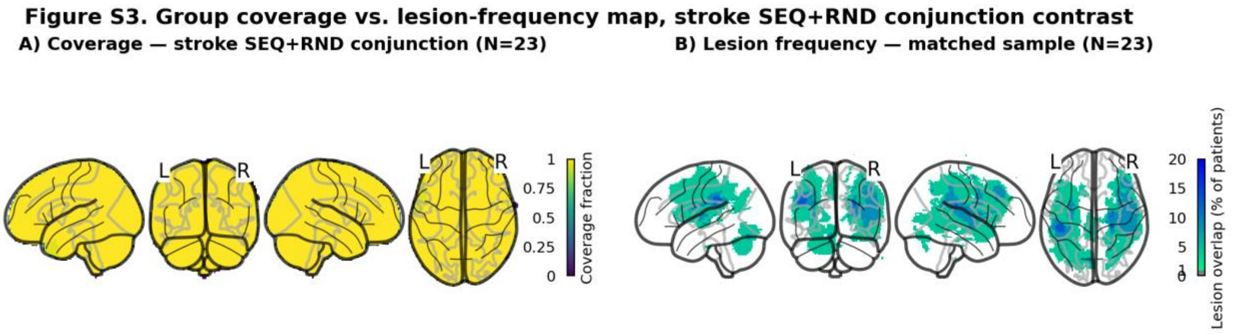

***Figure S6.*** *Group coverage vs. lesion-frequency map for the stroke cohort's SEQ+RND (all-runs) conjunction contrast. A) Contrast-specific coverage fraction, the proportion of the N=23 stroke participants entered into this second-level model with valid, non-lesioned signal at each voxel. B) Lesion-frequency map (FSL/lesionnorm pipeline, matched to the same N=23 sample), the percentage of participants with lesion involvement at each voxel, shown at the same MNI extent for direct visual comparison. Four-view glass-brain projections (sagittal left, coronal, sagittal right, axial), MNI152 space.*

#### S7. Extended ROI results

Full statistics for the a priori ROI mixed-effects models (main manuscript Results, Region of interest (ROI) analysis) are summarized below for the 11 primary learning-network ROIs (pre-SMA, bilateral PMd, bilateral PMv, bilateral cerebellum lobule VI, bilateral caudate, bilateral putamen) across the three tested fixed effects (Group, Stage, Group × Stage), with Benjamini–Hochberg FDR correction applied within each effect family (N=23 stroke, 14 control; consistent across the ≥25% and ≥50% ROI coverage thresholds and, with the exception of the pre-SMA Stage effect, across the sub-LRN010 exclusion – Supplementary Results S10). Only the 8 FDR-significant ROI × effect combinations are tabulated below; the exhaustive 17-ROI × all-contrasts table is not reproduced here.

##### Table S7: Region-of-interest model statistics (FDR-significant effects)

| ROI | EFFECT | F (DF1, DF2) | P | Q_FDR |
|---|---|---|---|---|
| **PUTAMEN (L)** | Group | 9.12 (1, 37) | .005 | .031 |
| **PUTAMEN (R)** | Group | 8.62 (1, 37) | .006 | .031 |
| **CEREBELLUM VI (R)** | Stage | 10.34 (1, 111) | .002 | .018 |
| **CEREBELLUM VI (L)** | Stage | 9.07 (1, 111) | .003 | .018 |
| **PMV (L)** | Stage | 6.59 (1, 108) | .012 | .039 |
| **CAUDATE (R)** | Stage | 6.21 (1, 111) | .014 | .039 |
| **PUTAMEN (R)** | Stage | 5.59 (1, 111) | .020 | .044 |
| **CAUDATE (L)** | Stage | 5.24 (1, 111) | .024 | .044 |

#### S8. Functional connectivity (gPPI) results

*Preregistered primary M1–M1 connectivity results.* All three PRIMARY seed-based gPPI tests (bilateral M1 hand-knob seed pair, tested separately as ipsi- and contralesional relative to each participant's affected hand; N=22 stroke, because one stroke participant with bilateral/multifocal lesions and no single defined ipsilesional hemisphere was excluded from all lateralized-seed connectivity analyses, Supplementary Methods) are summarized here against voxel-wise FWE correction ($p<.05$, whole-brain unless noted). Learning-related change (SEQ Early-minus-Late, RND Early-minus-Late, and the PRIMARY SEQ−RND Early-minus-Late contrast, both directions, both seeds) was null at voxel-FWE for all 12 tested combinations; the strongest uncorrected exploratory peak was Z=6.53 (contralesional seed, RND Early-minus-Late, MNI [-10.5,61.5,31.5]). The behavioral covariate test (SEQ−RND Early-minus-Late × winsorized performance-convergence slope, the PRIMARY contrast) was likewise null at voxel-FWE in both seeds and covariate directions (0/6 combinations, consistent with Table S8). One non-PRIMARY covariate contrast, ipsilesional seed, RND Early-minus-Late, positive covariate direction, reached voxel-FWE at a single voxel (Z=7.23, MNI [53.5,-18.5,41.5]); because this falls outside the pre-specified SEQ−RND PRIMARY contrast, it is reported as a secondary/exploratory observation rather than a confirmatory finding. The lesion-load moderation test (lag × corrected whole-brain lesion-load-fraction interaction) was the one PRIMARY gPPI family with a voxel-FWE-surviving PRIMARY-contrast result: contralesional seed, SEQ−RND Early-minus-Late, negative-interaction direction, 1 cluster/1 voxel, Z=7.85, MNI [-2.5,-10.5,-12.5], matching Table S8, row 5. As in Table S8, this isolated finding should be interpreted cautiously: it reflects a single voxel and has not been corroborated by an independent stability check. The ipsilesional-seed lesion-load moderation contrast was null in both directions across all three task contrasts.

*Exploratory connectivity-seed analyses.* The six secondary/exploratory seed-variants (pre-SMA, midline; putamen, ipsi- and contralesional; SPL/IPS, ipsi- and contralesional; dACC/CMA, midline; Supplementary Methods) were tested for the same three learning-related-change contrasts as the PRIMARY M1–M1 pair. None reached voxel-FWE correction in the stroke cohort — 0/36 tested seed-variant × contrast × direction combinations. Exploratory uncorrected peaks ($p<.001$, $k\geq10$) were present in most combinations, generally in occipital, parietal, or contralateral sensorimotor regions rather than a consistent network, e.g., ipsilesional SPL/IPS showed a 375-voxel cluster for RND Early-minus-Late (Z=6.12, MNI [-4.5,-82.5,-0.5]), and ipsilesional putamen showed a 151-voxel cluster for the same contrast (Z=7.03, MNI [19.5,-10.5,-2.5]). As none survive correction, these are reported as descriptive only and are not interpreted as evidence of learning-related connectivity change involving these seeds.

*Small-volume-corrected gPPI results.* To increase sensitivity within each seed's own anatomical territory, the PRIMARY M1–M1 and secondary putamen learning-related-change contrasts were additionally tested with small-volume correction (SVC; bilateral M1 hand-knob mask for the M1 seeds, bilateral putamen mask for the putamen seeds) rather than whole-brain FWE. All SVC results were null in the stroke cohort: 0/12 tested combinations (2 seeds × 3 contrasts × 2 directions) reached voxel-FWE-SVC for the M1 seeds, and 0/12 for the putamen seeds; the same pattern held in controls. Uncorrected exploratory activity within the M1 SVC mask was minimal to absent (0 clusters at $p<.001$, $k\geq3$, in every M1 SVC test, stroke and controls).

**S9. Lesion-load moderation results (activation)**

*Full interaction-contrast results.* The pre-specified lag × lesion-load interaction was tested across the three task contrasts (SEQ, RND, SEQ−RND Early-minus-Late), both interaction directions (positive, negative), the corrected and raw lesion-load metrics, and the primary 6-mm and sensitivity 8-mm group-level smoothing arms (24 unique combinations, N=23 stroke). At the primary 6-mm smoothing level, every combination was null at voxel-wise FWE correction ($p<.05$), for both metrics; the strongest uncorrected exploratory peak was the PRIMARY SEQ−RND contrast's negative-interaction direction (corrected metric: 41 clusters, 1415 voxels, Z=6.60, MNI [-36.5,-36.5,25.5]; raw metric nearly identical, Z=6.61, same location). Under the 8-mm group-level smoothing sensitivity arm, one combination reached voxel-FWE: the SEQ−RND Early-minus-Late negative-interaction contrast, corrected metric, 1 cluster/1 voxel, Z=6.25, MNI [-0.5,-62.5,-12.5], consistent with the corresponding result reported in Table S8 (reported there as Z≈6.25). All other 8-mm combinations, and both directions of the SEQ-only and RND-only contrasts at 8-mm, were null.

*Corrected vs. raw metric comparison.* Because the whole-brain lesion-load metric was control-baseline-corrected by default (see below), the raw (uncorrected) metric was retained throughout as a SENSITIVITY comparison, per Table S1's analysis pipeline. The one voxel-FWE-surviving combination above is robust to this choice: under the raw metric, the same contrast and direction (8-mm, SEQ−RND Early-minus-Late, negative interaction) also reached voxel-FWE, at an almost identical location and magnitude (1 cluster/1 voxel, Z=6.24, MNI [-0.5,-62.5,-12.5]). Every other combination that was null under the corrected metric was also null under the raw metric, at both smoothing levels. The 8-mm finding is therefore not an artifact of the control-baseline correction step, though it remains, as stated in Table S8, a single-voxel result confined to one non-primary smoothing arm and should be interpreted accordingly.

*Control-baseline correction audit.* The correction (Supplementary Methods) subtracts the control cohort's median raw whole-brain non-brain-tissue fraction from each stroke participant's own raw fraction, floored at zero. The control median was 0.0955 (n=14 controls, raw fraction range 0.0939–0.1061), confirming the raw metric's earlier characterization as dominated by a generic, lesion-independent component rather than lesion burden specifically. Applied to the stroke cohort (n=23, raw fraction range 0.0938–0.1769), five participants' corrected values floored to zero (raw fraction at or below the control median), and the remainder ranged up to 0.0814 (the highest-lesion-load participant). This correction produced a non-negative lesion-load distribution: a floor at zero for low-burden participants and a graded range for the rest, consistent with the correction functioning as intended.

**S10. Summary of preregistered confirmatory brain–behavior analyses**

Table S8 summarizes the five preregistered confirmatory tests of brain–behavior association reported in the main manuscript (Results, Brain–behavior and connectivity analyses). Analyses using the non-winsorized performance-convergence slope and additional exploratory analyses (small-volume-corrected ROIs, additional gPPI seeds, and alternative smoothing and early–late parameterizations) are reported in Sections S8 and S12.

##### Table S8: Summary of preregistered confirmatory brain–behavior analyses

| ANALYSIS FAMILY | PRE-SPECIFIED CONTRAST | N (STROKE) | OUTCOME (VOXEL-FWE, P<.05) | NOTE |
|---|---|---|---|---|
| **ACTIVATION GLM — BIVARIATE LAG COVARIATE** | SEQ−RND Early-minus-Late, winsorized behavioral covariate | 23 | Null (0 clusters) | Strongest exploratory peak Z=5.98 (uncorrected) |
| **ACTIVATION GLM — LESION-LOAD MODERATION** | Lag × corrected lesion-load interaction | 23 | Null (0/24 tested combinations) | 8-mm smoothing arm: single-voxel interaction Z≈6.25, confined to one non-primary smoothing arm (Section S12) |
| **GPPI — M1–M1 LEARNING-RELATED CHANGE** | SEQ/RND Early-minus-Late, ipsi- + contralesional seeds | 22 | Null (0/24 tested combinations) | — |
| **GPPI — M1–M1 BEHAVIORAL COVARIATE** | SEQ−RND Early-minus-Late × lag | 22 | Null (0/6 tested combinations) | — |
| **GPPI — M1–M1 LESION-LOAD MODERATION** | Lag × corrected lesion-load interaction, contralesional seed | 22 | 1 voxel FWE, Z=7.85, MNI[−2.5,−10.5,−12.5] | Not reproduced using the ipsilesional seed; considered exploratory (Section S8) |

FWE = voxel-wise family-wise error correction ($p < .05$). Reported effect-size/precision values for null results correspond to exploratory (uncorrected) peak statistics, provided to inform the design and power calculations of future studies; they should not be interpreted as confirmatory effect estimates.

#### Sensitivity Analyses

##### S11. ROI robustness checks

Two aspects of the extended ROI results (Section S7) were tested for robustness: the minimum ROI-coverage threshold used to exclude low-coverage subject×ROI cells (Section S5), and the inclusion of sub-LRN010 (excluded from all other whole-brain imaging analyses for failed spatial normalization; see Methods). Table S9 summarizes both checks.

#### Table S9: ROI coverage-threshold and sample sensitivity checks

| ANALYSIS | PRIMARY RESULT | SENSITIVITY RESULT | INTERPRETATION |
|---|---|---|---|
| **COVERAGE THRESHOLD (≥25% VS. ≥50% MINIMUM ROI COVERAGE)** | 9/11 ROI×effect combinations FDR-significant at the pre-specified ≥25% threshold (N=24 stroke, 14 control) | Identical set of combinations FDR-significant at the stricter ≥50% threshold; only trivial shifts in F-statistics and denominator df | Findings are robust to the specific coverage threshold used |
| **SUB-LRN010 EXCLUSION (N=24 VS. N=23 STROKE)** | 9 ROI×effect combinations FDR-significant at N=24 | 8/9 remained FDR-significant at N=23 with comparable effect sizes (e.g., putamen (L) Group: F=9.25→9.12, q=.025→.031); the pre-SMA Stage effect, already the weakest of the nine, fell below threshold (p=.045–.051, q=.071–.080) | Findings are robust to the sub-LRN010 exclusion except the pre-SMA Stage effect, which should be interpreted as more marginal than the other eight |
| **RIGHT PMD GROUP EFFECT (EXPLORATORY)** | p=.053 (N=24), not FDR-significant | p=.025 (N=23), still not FDR-significant (q=.069) | No new FDR-significant findings emerged under either coverage threshold or sample |

##### S12. Activation robustness and sensitivity checks

Two aspects of the activation lesion-load moderation and whole-brain learning-contrast results (Table S8; Section S6) were tested for sensitivity to analysis parameterization: use of the raw, non-winsorized behavioral covariate in place of the winsorized PRIMARY covariate, and an additional 8-mm group-level smoothing pass on top of the 6-mm first-level smoothing used throughout. Table S10 summarizes both checks.

#### Table S10: Activation covariate and smoothing sensitivity checks

| ANALYSIS | PRIMARY RESULT | SENSITIVITY RESULT | INTERPRETATION |
|---|---|---|---|
| **LAG-COVARIATE MODEL, ALTERNATIVE REPARAMETERIZATIONS (RAW, NON-WINSORIZED COVARIATE)** | Winsorized-covariate PRIMARY model (Table S8): null at whole-brain FWE across all three parameterizations | Raw covariate reached FWE at a small number of voxels in all three re-analyses, but at a different peak location and cluster topology each time (1-run window/6-mm: 6 clusters/16 voxels, Z=11.38, MNI [53.5,-28.5,5.5]; 2-run-block window/6-mm: 2 clusters/23 voxels, Z=8.50, MNI [29.5,29.5,31.5]; 1-run window/8-mm: 3 | Raw-covariate results are unstable across reparameterizations and are not interpreted as a robust effect |

| | | | |
|---|---|---|---|
| | | clusters/67 voxels, Z=6.41, MNI [29.5,27.5,27.5]) | |
| **WHOLE-BRAIN LEARNING CONTRASTS, 6-MM VS. 8-MM GROUP-LEVEL SMOOTHING (SECTION S6)** | Null at 6-mm in both cohorts | Remained null in stroke at 8-mm; one additional cluster reached FWE in controls at 8-mm only (SEQ Early-minus-Late, negative direction, 2 voxels, Z=11.21, MNI [-4.5,27.5,5.5]) | Small, smoothing-dependent cluster confined to controls; not interpreted as a reversal of the null learning-contrast pattern (Section S6) |
| **LESION-LOAD MODERATION MODEL, 6-MM VS. 8-MM SMOOTHING (TABLE S8)** | Null at 6-mm for both the corrected and raw lesion-load metric | Single FWE-significant voxel at 8-mm for both the corrected (Z=6.25, MNI [-0.5,-62.5,-12.5]) and raw (Z=6.24, same location) metric | The 8-mm result is not driven by the control-baseline correction, but remains an unconfirmed, smoothing-dependent finding confined to one non-primary smoothing arm, consistent with Table S8's designation |
